# The XPF-like domain in SHOC1 required for homologous recombination and safeguarding autosome from meiotic silencing of unsynapsed chromatin

**DOI:** 10.1101/2025.05.28.656576

**Authors:** Yuxiang Zhang, Qian Sun, Zhiyong Ji, Ren Mo, Yuchuan Zhou, Jingpeng Zhao, Shuai Xu, Na Li, Yifan Sun, Haowei Bai, Erlei Zhi, Sha Han, Huixing Chen, Jing Zhang, Dewei Qian, Xinjie Bu, Yuhua Huang, Ruhui Tian, Ying Guo, Jinxing Lv, Liangyu Zhao, Chao Yang, Fujun Zhao, Peng Li, Zhi Zhou, Zheng Li, Chencheng Yao

## Abstract

During meiosis, a group of evolutionarily conserved ZMM proteins plays essential roles in stabilizing the recombination intermediates and promoting crossover (CO) formation. In mice, SHOC1 forms a trimeric complex with the other two ZMM proteins, SPO16 and TEX11, to bind recombination intermediates after strand invasion. Although genetic variants of *SHOC1* are clinically associated with meiotic arrest and male infertility, their precise molecular mechanisms and evolutionarily conserved functions in human gametogenesis remain enigmatic. Here, we delineated species-specific divergences between human and mouse SHOC1 complex, and identified a missense variant within the XPF-like domain in *SHOC1* (c.A1769G:p.Q590R) that was associated with meiotic arrest and non-obstructive azoospermia (NOA). The disorder of the XPF-like domain in SHOC1 impaired DNA double-strand breaks repair by compromising its ability to bind branched DNA structures and the recruitment of M1AP, REDIC1, and ZMM factors to recombination intermediates, ultimately abolishing CO formation. Furthermore, the variant disrupted dynamic 3D chromatin structure in pachytene spermatocytes and induced defects in homologous chromosome synapsis. More importantly, the XPF-like domain in SHOC1 was revealed to prevent autosome intrusion into the sex body compartment, thereby safeguarding critical autosomal loci from meiotic silencing of unsynapsed chromatin (MSUC). Overall, our study demonstrated that the XPF-like domain in SHOC1 is required for homologous recombination and safeguarding autosome from MSUC in meiosis.

## INTRODUCTION

Infertility affects an estimated 15% of couples of reproductive ages globally, presenting a significant medical and societal challenge(*1*). Male infertility accounts for approximately 50% of these cases, with non-obstructive azoospermia (NOA) being the most severe type(*2*). Meiotic arrest, a subtype of NOA, is widely known to have multiple genetic origins, including chromosome abnormalities, Y chromosome microdeletions, and monogenic variants(*3, 4*). However, the pathogenic mechanisms of genetic disorder derived meiotic arrest remain elusive.

Meiosis is essential for the production of haploid gametes in mammals, during which homologs undergo pairing, synapsis, and recombination. Meiotic recombination is orchestrated through a sophisticated sequence of events. Programmed DNA double-strand breaks (DSBs) are formed by the SPO11 enzyme during the early stages of meiotic prophase I and are quickly processed to generate 3’ single-stranded DNA (3’-ssDNA) overhangs(*5, 6*). These overhangs are subsequently enveloped by the single-strand DNA binding protein complex, replication protein A (RPA) and the RAD51/DMC1 recombinase complex. The RPA complex collaborates with meiosis-specific proteins such as MEIOB and SPATA22 to facilitate the recombination process(*5-7*). Nascent displacement loops (D-loops) migrate towards the direction of repair synthesis. One resected DSB end can be cross-linked with a partner duplex in a stable, and discrete state namely single-end invasion (SEI). In yeast, the SEI intermediates rely on the ZMM factors, including Zip1, Zip2, Zip3, Zip4, Msh4-Msh5 (the MutSγ complex), Mer3, and Spo16, to stabilize nascent joint molecules and coordinately promote polymerization of synaptonemal complex (SC)(*8, 9*). These recombination intermediates are bound by RNF212 and HEI10 (an ortholog of Zip3) to regulate their stability and further recruit CDK2 and MLH1-MLH3, eventually repaired into class L crossovers (COs)(*10, 11*).

Three ZMM proteins, Zip2, Zip4, and Spo16, form a functionally conserved protein complex known as “ZZS”, which was initially described in yeast(*12*). In mice, all three ZZS proteins localize to chromosome axes as discrete foci and have similar foci kinetics in meiotic prophase I(*13*). Notably, mouse Shortage in chiasmata 1 (SHOC1), orthologs of yeast Zip2 has been reported to play indispensable roles in spermatogenesis(*14-16*). During meiosis, SHOC1 is recruited to D-loops following strand invasion, where it binds specific DNA to catalytically stabilize recombination intermediates and drive DNA synthesis. Also, it recruits SPO16 to maintain the stabilization of SHOC1 and proper localization of TEX11 (ortholog of Zip4) and MSH4(*13, 14, 17*). MSH4 and MSH5 form a clamp-like heterodimer to stabilize the DNA structure associated with strand invasion. Also, they are responsible for the recruitment of MLH1 and MLH3 to the COs sites in pachytene(*18, 19*). *Shoc1*^-/-^ and *Spo16*^-/-^ mice showed severe defects in synapsis and meiotic arrest at zygotene-like or early pachytene-like stage without CO formation(*13, 14*), while knockout of *Tex11* in mice caused a milder defect in synapsis and reduced CO formation, leading to meiotic metaphase I (MMI) arrest(*20, 21*). Recently, two novel proteins M1AP and REDIC1, have been characterized to have similar locations and functions to the ZZS proteins in facilitating CO formation and meiotic progression in males(*22, 23*).

The XPF-like domain, which is indispensable for binding with DNA structures, is conserved in yeast Zip2, mouse SHOC1 as well as human C9orf84 (also referred to SHOC1). The XPF-like proteins, including MUS81, EME1, FANCM, and SHOC1, are distinguished by helicase domain, ERCC4-like nuclease domain, and conserved Ercc4-helix-hairpin-helix (HhH)L cores(*15*). They are proved to be fundamental in meiotic recombination typically functioning as heterodimers, including XPF-ERCC1 complex, MUS81-EME1/2 complexes, FANCM-FAAP24 complex, and SHOC1-SPO16 complex(*13*). Specifically, Mus81-Eme1 complexes are revealed to be required for resolution of double Holliday junctions into class II COs(*24*). Also, FANCM-FAAP24 complexes are showed to promote class I interfering COs and suppress class II non-interfering COs(*25*). In mice, SHOC1-SPO16 complex cooperates with MSH4-MSH5 to recognize and stabilize early recombination intermediates, recruit downstream CO factors to promote the formation of class L COs, and coordinate the formation of COs with assembly of the SC(*13*). It has been demonstrated that recombinant XPF-like domain in human SHOC1 preferentially binds branched DNA but lacks detectable endonuclease activity *in vitro(17)*. However, the *in vivo* roles and critical residues of the XPF-like domain in SHOC1 during meiosis still remain unclear.

Herein, we revealed the differences between human and mouse SHOC1 complexes and identified a missense variant within the XPF-like domain in *SHOC1* (c.A1769G:p.Q590R) that was associated with non-obstructive azoospermia (NOA). Intriguingly, the knock-in (KI) mice mimicking the patient’s variants showed defects in DSBs recombination and homologous chromosome synapsis. Furthermore, abnormal silencing of unsynapsed autosomes in sex body took place in pachytene spermatocytes of *Shoc1* KI mice. Overall, our study offers novel insights into meiotic recombination and provides a prospective molecular target for the clinical diagnosis and treatment of infertility.

## RESULTS

### Human SHOC1 interacted with M1AP, REDIC1, and ZZS protein TEX11, but not C1orf146 (an ortholog of SPO16)

To determine the components of SHOC1 complex in human, we performed co-immunoprecipitation (co-IP) assay. Human FLAG-tagged SHOC1 was co-expressed with human HA-tagged M1AP or REDIC1, or MYC-tagged TEX11 or C1orf146 in HEK293T cells. Proteins were immunoprecipitated from cell lysates by tag-specific antibodies. A subsequent Western blot (WB) showed that human SHOC1 bond specifically with M1AP, REDIC1 and ZZS protein TEX11. However, we did not detect a specific interaction between human SHOC1 and C1orf146, which was different from the previous report in mice(*13*) (Figure 1A,B). Specifically, mouse SHOC1 could interact with all partners including TEX11, M1AP, REDIC1, and SPO16. Also, the human canonical “ZZS” complex were further confirmed by bimolecular fluorescence complementation (Bi-FC) assay, which showed that the Venus signals could be detected in SHOC1 and TEX11 co-expressing cells instead of SHOC1 and C1orf146, or TEX11 and C1orf146 co-expressing groups (Figure 1C). Thus, we proposed distinct models for the SHOC1 complex in human and mice (Figure 1D).

**Figure 1.**
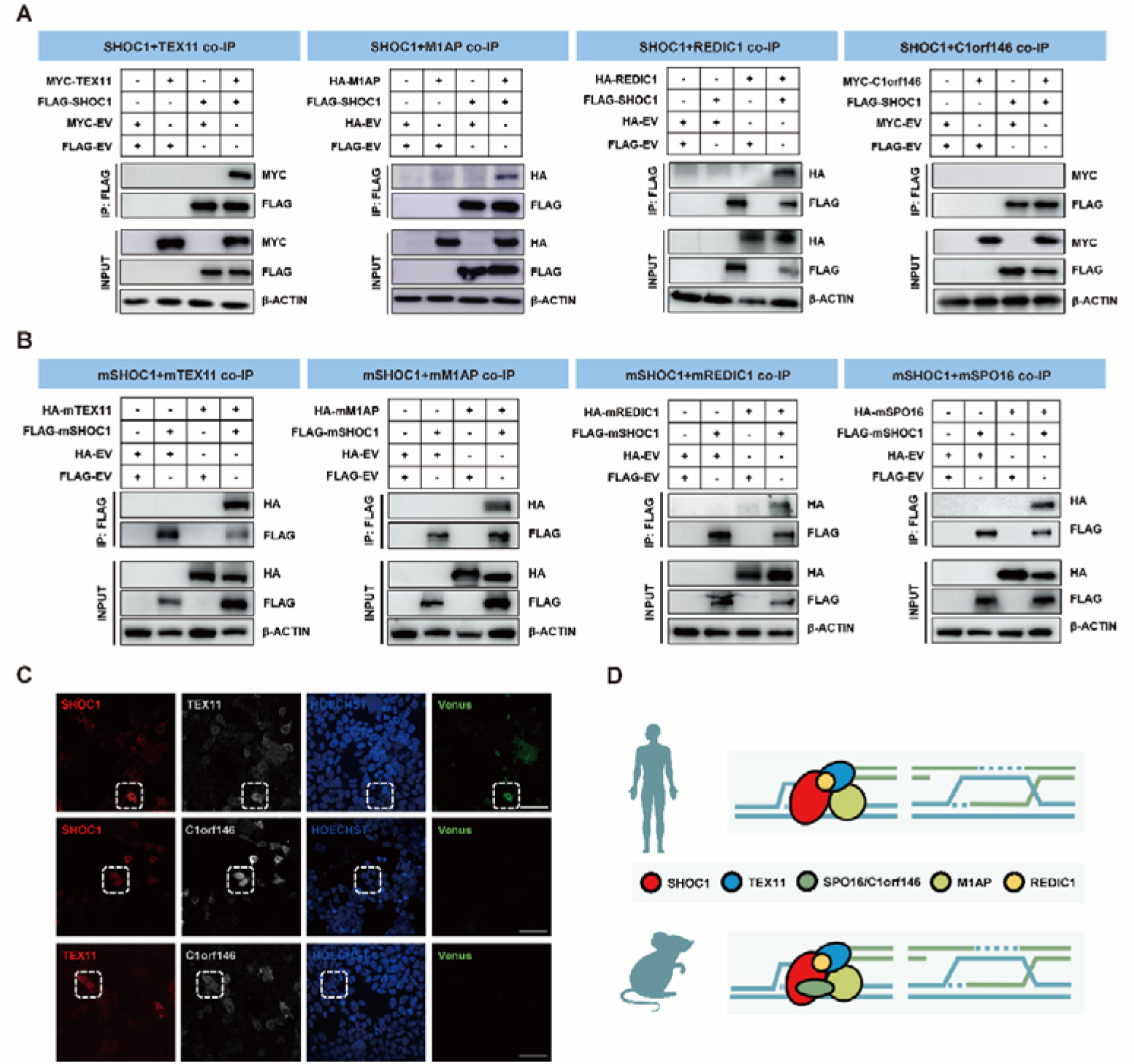
Distinct models for the SHOC1 complex in human and mouse. **(A)** Co-IP demonstrated the interaction of human SHOC1 (detected via N-terminal FLAG tag) with M1AP, REDIC1, and ZZS protein TEX11, but not with C1orf146 (detected via N-terminal HA or MYC tag). Input lysates were included. **(B)** Co-IP confirmed the interaction of mouse SHOC1 (mSHOC1) (detected via N-terminal FLAG tag) with mM1AP, mREDIC1, and ZZS proteins mTEX11 and mSPO16 (detected via N-terminal HA tag). Input lysates were included. **(C)** Bi-FC revealed Venus fluorescence signals in human SHOC1- and TEX11-coexpressing cells, but not in SHOC1/C1orf146-or TEX11/C1orf146-coexpressing groups. Scale bars, 50 μm. **(D)** Schematic illustration of the SHOC1 complex components in human and mouse.

To further determine the specific region of SHOC1 involved in interactions with other partners, we co-expressed M1AP, REDIC1 or TEX11 full-length proteins and various SHOC1 truncated proteins (Human: N-terminal 1-576, C-terminal 577-1444, ΔF1 aa1-288del, ΔF2 aa289-576del, ΔF3 aa577-1098del, ΔF4 aa1099-1444del; Mouse: N-terminal 1-625, C-terminal 626-1482, ΔF1 aa1-312del, ΔF2 aa313-625del, ΔF3 aa626-1154del, ΔF4 aa1155-1482del) in HEK293T cells (Figure S1A,B). The co-IP and subsequent WB analysis demonstrated that truncation of specific regions (ΔF1-ΔF4) of SHOC1 did not disrupt the binding of M1AP or REDIC1 (Figure S1C,D), suggesting that SHOC1 has multiple interaction sites with M1AP or REDIC1. Notably, removal of the N-terminal region of SHOC1 completely abolished TEX11 binding in both mouse and human, suggesting that the isolated SHOC1 N-terminal region is essential for interaction with TEX11. Interestingly, ΔF1 and ΔF2 fragments of mouse SHOC1 did not affect TEX11 binding, indicating the presence of TEX11 binding sites in both fragments. In contrast, ΔF1 and ΔF2 fragment in human SHOC1 completely abolished the TEX11 binding, underscoring the necessity of an intact N-terminal region for maintaining the SHOC1-TEX11 interaction (Figure S1C,D). These findings further suggested that there are differences in the interaction patterns between human and mouse SHOC1 and its partners, particularly with TEX11, indicating a potential species-specific function of human SHOC1 in meiosis that warrants further investigation.

### Missense variant within the XPF-like domain in *SHOC1* (p.Q590R) was associated with meiotic arrest and NOA

Idiopathic NOA caused by meiotic arrest is a common cause of male infertility and has many genetic origins. Here, we screened for potential variants in a cohort of 171 patients with meiotic arrest and found bi-allelic *SHOC1* variants were associated with NOA. In addition to the compound heterozygous variants in *SHOC1* (M3: c.C1582T:p.R528X and M4: c.231_232del:p.L78Sfs*9), homozygous loss-of-function (LoF) variants (M5: c.1194delA:p.L400Cfs*7 and M6: c.1347delT:p.D450Tfs*13) in our previous work(*26*), novel compound heterozygous variants (M1: c.A1769G:p.Q590R and M2: c.416_419del:p.S139SFs*9) in family 3 and a homozygous splicing variant (M7: c.G2738-1A) of *SHOC1* in family 4 were identified after the genetic analyses pipeline in the current study. All variants were absent or had low allele frequencies in public databases. Sanger sequencing validated all above *SHOC1* variants in pedigrees affected by NOA, consistent with autosomal recessive inheritance patterns following pedigree co-segregation analysis (Figure 2). Clinical details for each patient are outlined in supplementary table S3.

**Figure 2.**
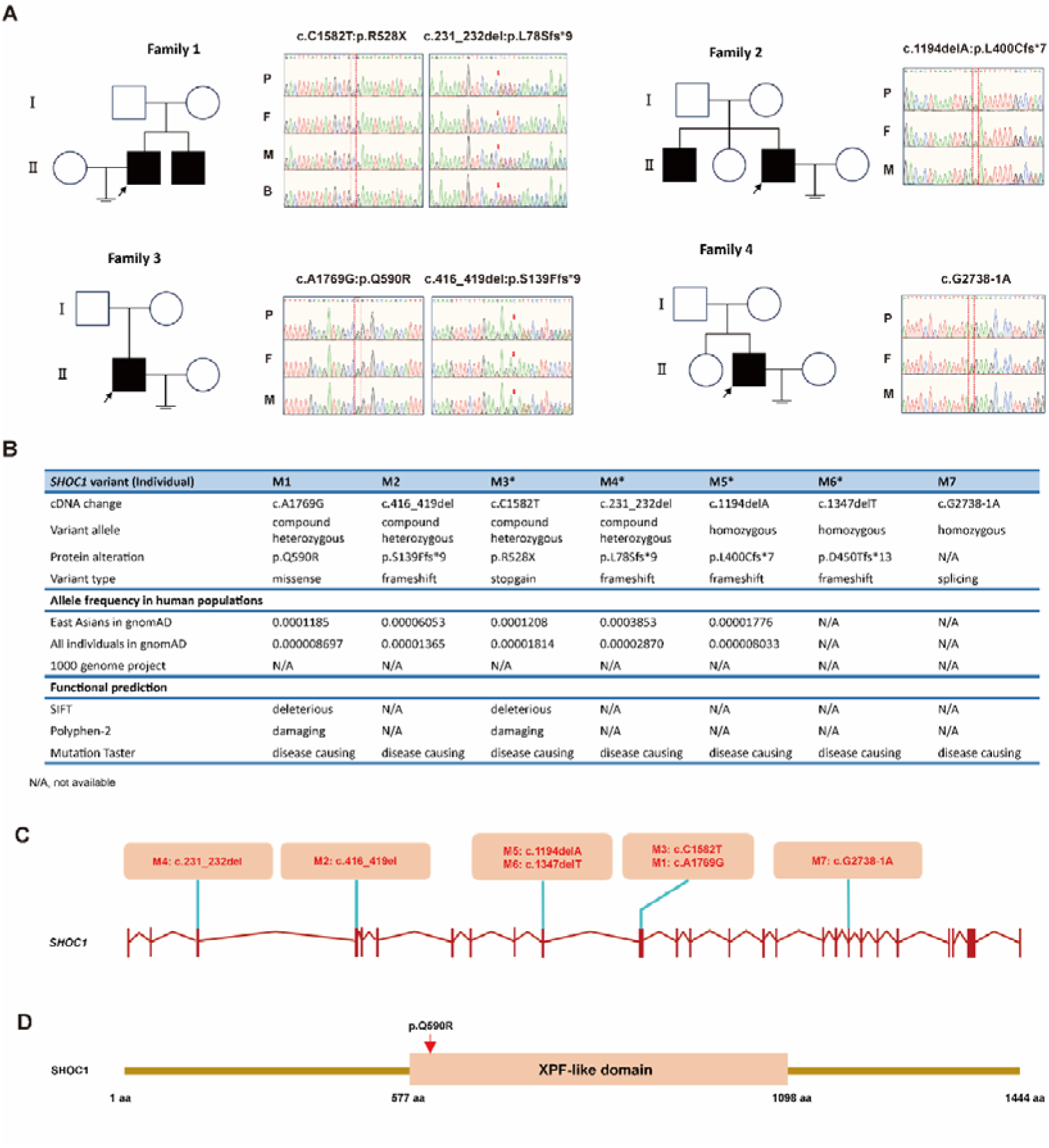
Bi-allelic *SHOC1* variants were identified in NOA-affected patients with meiotic arrest. **(A)** Pedigrees of infertile families carrying *SHOC1* variants. WES findings were validated by Sanger sequencing. Probands are indicated by black arrows, and variant-affected nucleotides are highlighted with red dash line boxes or arrows. **(B)** Summary of *SHOC1* variants identified in this study, with asterisks denoting variants reported in our previous publication. **(C)** The localization of identified *SHOC1* variant residues. **(D)** Schematic diagram of the human SHOC1 protein, annotated with functional domains. The p.Q590R variant is highlighted by a red arrow.

All identified *SHOC1* variants in this study included one missense (M1), four frameshifts (M2, M4, M5 and M6), one stop-gain (M3), and one splicing (M7) types. Among them, the variant M2 to M6 were predicted to result in the deletion of the majority of regions of SHOC1 protein and assumed as pathogenic using pathogenicity prediction tools including SIFT, PolyPhen-2 and MutationTaster (Figure 2B). Next, we transfected the full-length wild-type (WT) and mutant cDNA constructs of *SHOC1* to HEK293T cells. As a result, we found protein degradation in the variant (M4), and truncations in the variants (M2, M3, M5, and M6) (Figure S2A). Therefore, we proposed that all LoF variants (M2-M6) in *SHOC1* were potential disease-causing and associated with male infertility in the affected individuals. The variant (M7) was located in the region of the canonical splice donor site at the boundary of exon 21. To assess the impact of the variant on *SHOC1* splicing, minigene vectors containing the genomic sequence spanning exon 21 and flanking introns of the *SHOC1* gene were transfected into HEK293T, followed by RT-PCR. The WT SHOC1 minigene showed a canonical splicing pattern, while the splicing variant M7 resulted in a band smaller than that of WT (Figure S2B). Sanger sequencing showed that the splicing variant caused skipping of exon 21 (Figure S2C), which was assumed to produce truncated proteins deficient in functional segments within the XPF-like domain. This result suggested that the splicing variant M7 in *SHOC1* (c.G2738-1A) induced aberrant mRNA splicing (Figure S2D).

The only identified missense variant M1 in family 3, situated within the functional XPF-like domain, was also predicted to be pathogenic by all three tools (Figure 2B), underscoring its importance of this residue within the XPF-like domain in spermatogenesis. Testicular biopsies of the proband harboring the compound heterozygous variants (M1: c.A1769G:p.Q590R and M2: c.416_419del:p.S139Ffs*9) of *SHOC1* in family 3 were obtained and the hematoxylin and eosin (H&E) staining showed that all analyzed seminiferous tubules contained spermatogonia and spermatocytes but lacked spermatids and spermatozoa. In contrast, the testicular histopathology of testes in OA-affected patients showed normal spermatogenesis (Figure S3A). Moreover, IF staining and meiotic chromosomal spread analysis through various marker combination (SYCP3 & γ H2AX; DMC1 & PNA) also verified the meiotic arrest phenotype in this proband (Figure S3B-D). Collectively, these findings underscore a critical role of the XPF-like domain in SHOC1, especially the Q590 residue in human meiosis and establish one clinical-genetic paradigm for mechanistic dissection of male infertility caused by the missense variant M1 within the XPF-like domain (Figure S3E).

### *Shoc1* mutation (p.Q646R) resulted in sterility in both male and female mice

To determine the specific role of the XPF-like domain in SHOC1 during meiosis, we applied CRISPR/Cas9-mediated gene targeting to generate *Shoc1* mutant (p.Q646R) KI (*Shoc1* KI) mice mimicking the identified missense variant M1 (p.Q590R) within the XPF-like domain in *SHOC1* in family 3 (Figure 3A). Subsequent assessment of fertility in adult *Shoc1* KI mice revealed complete sterility in all tested homozygous males (Figure 3B). Furthermore, smaller ovaries and follicular dysplasia were found in adult *Shoc1* KI females (Figure S4). Additionally, adult KI male mice displayed significantly smaller testes and lower testis-to-body weight ratio compared to their littermate controls (Figure 3C,D). Computer-aided sperm analysis (CASA) showed the absence of sperm in the cauda epididymis of adult *Shoc1* KI mice (Figure 3E-G). Histological analysis demonstrated a complete lack of post-meiotic germ cells in all seminiferous tubules of the testes, with no mature sperm present in the cauda epididymis in adult *Shoc1* KI mice. In contrast, the control group exhibited abundant germ cells at various developmental stages in the testes and mature sperm in the cauda epididymis (Figure 3H,I). Notably, many MML spermatocytes with condensed nuclei and misaligned chromosomes were found in KI mice (Figure 3I).

**Figure 3.**
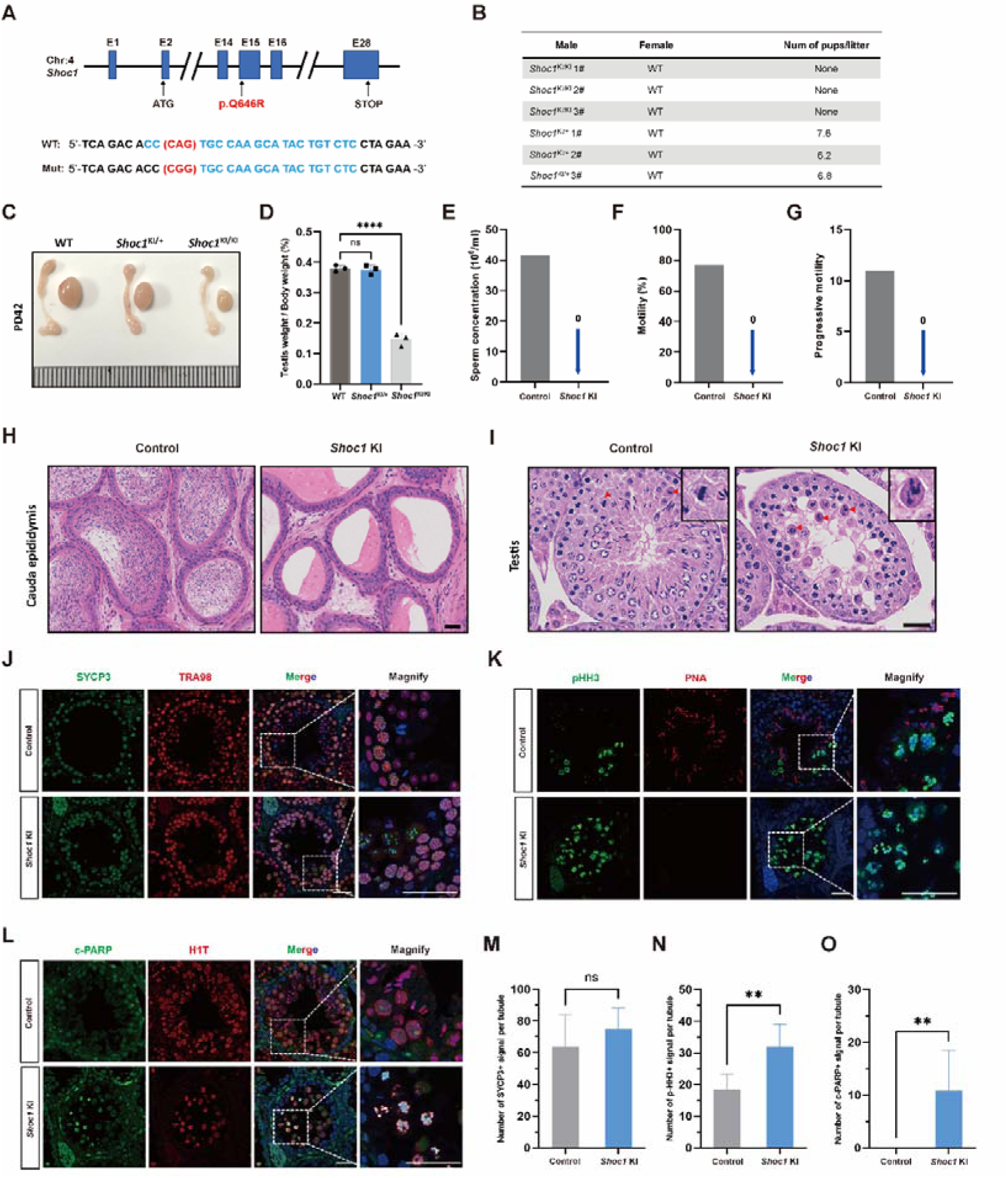
Male mice homozygous for the *Shoc1* missense variant (p.Q646R) exhibited spermatogenic failure and MMI arrest. **(A)** CRISPR/Cas9-mediated genome engineering strategy used to generate a C57BL/6J mouse model carrying the *Shoc1* mutation (p.Q646R). **(B)** Fertility assessment of *Shoc1* KI homozygous (*Shoc1*^KI/KI^) mice and littermate controls (*Shoc1*^KI/+^). **(C)** Representative image comparing testis size in adult *Shoc1* KI homozygous (*Shoc1*^KI/KI^) mice and littermate controls (*Shoc1*^KI/+^ and WT). **(D)** Testis-to-body weight ratios quantified for *Shoc1* KI homozygous (*Shoc1*^KI/KI^) mice and littermate controls (*Shoc1*^KI/+^ and WT) using one-way ANOVA; **** *P* < 0.0001; ns, not significant; error bars, mean ± SEM. **(E-G)** Epididymal sperm count and motility analyzed by CASA. **(H)** H&E-stained epididymal sections from adult *Shoc1* KI homozygous (*Shoc1*^KI/KI^) mice and littermate controls (*Shoc1*^KI/+^). Scale bars, 50 μm. **(I)** H&E-stained testicular sections from adult *Shoc1* KI homozygous (*Shoc1*^KI/KI^) mice and littermate controls (*Shoc1*^KI/+^). Scale bars, 50 μm. **(J)** IF staining of SYCP3 (green) and TRA98 (red) in testicular sections from adult *Shoc1* KI homozygous (*Shoc1*^KI/KI^) mice and littermate controls (*Shoc1*^KI/+^). Scale bars, 50 μm. **(K)** IF staining of pHH3 (green) and PNA (red) in testicular sections from adult *Shoc1* KI homozygous (*Shoc1*^KI/KI^) mice and littermate controls (*Shoc1*^KI/+^). Scale bars, 50 μm. (**L)** IF staining of c-PARP (green) and H1T (red) in testicular sections from adult *Shoc1* KI homozygous (*Shoc1*^KI/KI^) mice and littermate controls (*Shoc1*^KI/+^). Scale bars, 50 μm. (**M-O)** Quantification of SYCP3 (M), pHH3 (N), and c-PARP (O) positive signals in testicular sections from adult *Shoc1* KI homozygous (*Shoc1*^KI/KI^) mice and littermate controls (*Shoc1*^KI/+^) using two-tailed Student’s t-test; ** *P* < 0.01; ns, not significant; error bars, mean ± SEM.

Immunofluorescence (IF) analysis of testicular sections from adult male mice revealed spermatogenic failure in the KI group compared to controls. It was showed a significant reduction in TRA98-positive germ cells within the KI group (Figure 3J), while semi-quantitative assessment demonstrated that the number of SYCP3-positive spermatocytes remained comparable between groups (Figure 3J,M). Notably, the KI group exhibited complete depletion of post-meiotic haploid spermatids identified by PNA labeling, accompanied by a significant accumulation of pHH3-positive MMI spermatocytes (Figure 3K,N). Elevated number of c-PARP signals indicated enhanced apoptosis in KI seminiferous tubules, with H1T co-staining confirming apoptotic events specifically in arrested MMI spermatocytes (Figure 3L,O). These findings collectively demonstrated that *Shoc1* KI induced meiotic arrest at MMI, characterized by failure of meiotic progression, and subsequent apoptosis of arrested spermatocytes.

To clarify the exact substage of meiotic defects that occurred in *Shoc1* KI males, we further examined the progression of meiotic prophase I by immunostaining spermatocyte spreads for SYCP3 and γH2AX (a marker of DNA breaks). While leptotene, zygotene, pachytene, and diplotene spermatocytes were observed in both control and KI mice, a distinct population of cells (representing 41.9%) resembling pachytene spermatocytes but displaying aberrant γH2AX signals on autosomes (referred to as pachytene-like spermatocytes in this study) was exclusively identified in KI mice (Figure S5). Taken together, *Shoc1* KI male mice recapitulated the meiotic defects of the NOA-affected patient and further demonstrated the conserved glutamine residue (Q590 in human, Q646 in mice) within the XPF-like domain as indispensable for meiosis.

### The Q646 residue within the XPF-like domain in SHOC1 is required for CO formation by orchestrating SHOC1 complex assembly during meiotic recombination

MMI spermatocytes exhibiting chromosome misalignment often arise from defective CO formation between homologous chromosomes during prophase I(*27*). Thus, we quantified the number of MLH1 foci in pachytene stage, which mark sites of class I COs, in both KI and control groups. Control spermatocytes displayed 23.0 ± 1.5 MLH1 foci per nucleus, whereas the KI group showed nearly complete absence, with only 0.1 ± 0.3 foci per nucleus (Figure 4A,B). The HEI10 E3 ligase is another late recombination nodule component and is directed by MutLγ to stably accumulate at designated CO sites. Correspondingly, the number of HEI10 foci was greatly reduced in *Shoc1* KI mice (0.3 ± 0.7 vs. 22.8 ± 3.9) (Figure 4C,D). These findings confirmed that the p.Q646R mutation disrupted CO formation in KI groups.

**Figure 4.**
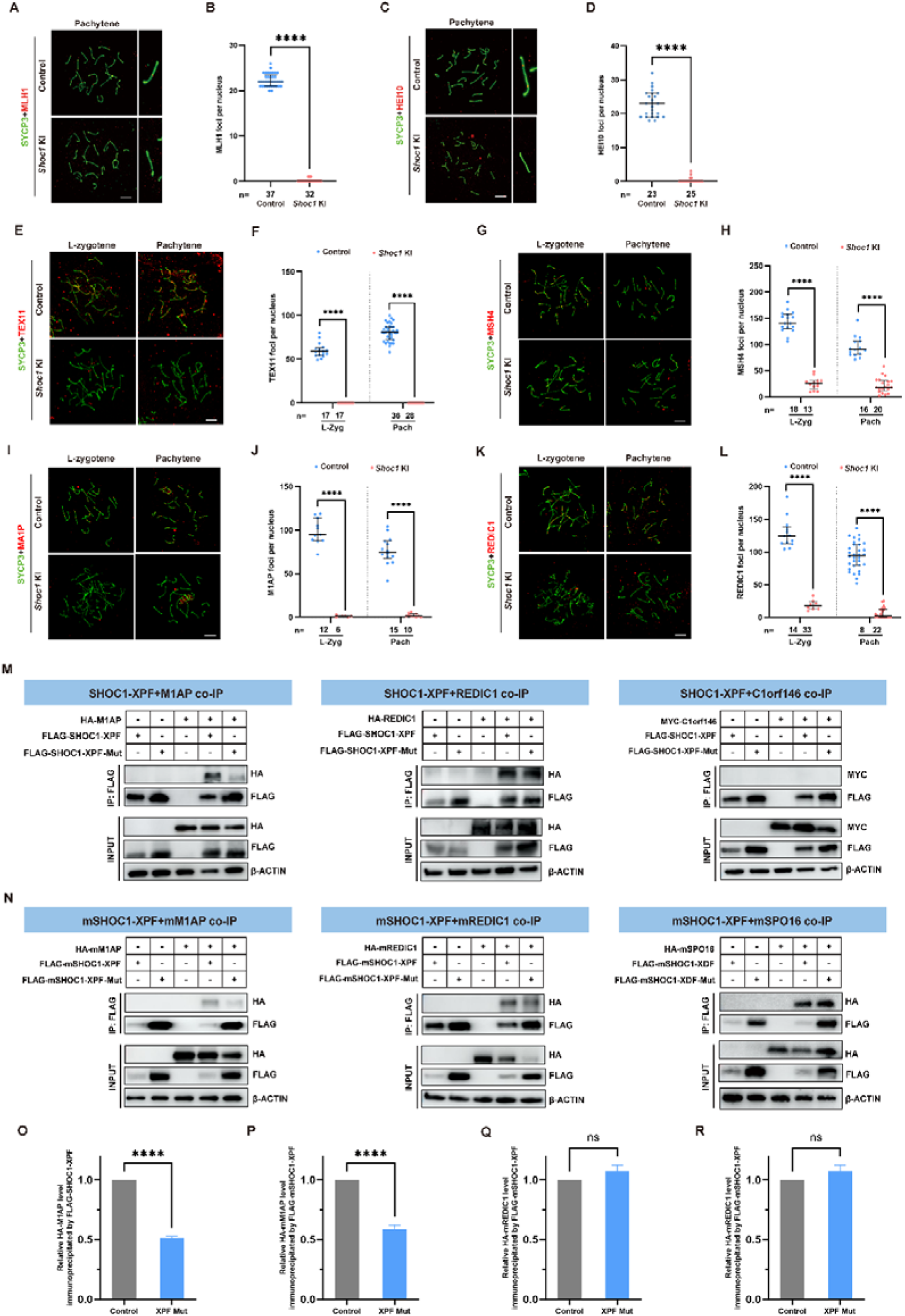
*Shoc1* KI mice exhibited defective CO formation and disrupted dynamics of meiotic recombination intermediates. **(A)** Representative images of spread spermatocytes from adult *Shoc1* KI homozygous (*Shoc1*^KI/KI^) mice and littermate controls (*Shoc1*^KI/+^) co-stained with SYCP3 (green) and MLH1 (red). Scale bars, 10 μm. **(B)** Quantification of MLH1 foci per cell at indicated meiotic stages using two-tailed Student’s t-test; **** *P* < 0.0001; error bars, mean ± SEM; n, the total number of nuclei analyzed. **(C)** Representative images of spread spermatocytes from adult *Shoc1* KI homozygous (*Shoc1*^KI/KI^) mice and littermate controls (*Shoc1*^KI/+^) co-stained with SYCP3 (green) and HEI10 (red). Scale bars, 10 μm. **(D)** Quantification of HEI10 foci per cell at indicated meiotic stages using two-tailed Student’s t-test; **** *P* < 0.0001; error bars, mean ± SEM; n, the total number of nuclei analyzed. **(E)** Representative images of spread spermatocytes from adult *Shoc1* KI homozygous (*Shoc1*^KI/KI^) mice and littermate controls (*Shoc1*^KI/+^) co-stained with SYCP3 (green) and TEX11 (red). Scale bars, 10 μm. **(F)** Quantification of TEX11 foci per cell at indicated meiotic stages using two-tailed Student’s t-test; **** *P* < 0.0001; error bars, mean ± SEM; n, the total number of nuclei analyzed. **(G)** Representative images of spread spermatocytes from adult *Shoc1* KI homozygous (*Shoc1*^KI/KI^) mice and littermate controls (*Shoc1*^KI/+^) co-stained with SYCP3 (green) and MSH4 (red). Scale bars, 10 μm. **(H)** Quantification of MSH4 foci per cell at indicated meiotic stages using two-tailed Student’s t-test; **** *P* < 0.0001; error bars, mean ± SEM; n, the total number of nuclei analyzed. **(I)** Representative images of spread spermatocytes from adult *Shoc1* KI homozygous (*Shoc1*^KI/KI^) mice and littermate controls (*Shoc1*^KI/+^) co-stained with SYCP3 (green) and M1AP (red). Scale bars, 10 μm. **(J)** Quantification of M1AP foci per cell at indicated meiotic stages using two-tailed Student’s t-test; **** *P* < 0.0001; error bars, mean ± SEM; n, the total number of nuclei analyzed. **(K)** Representative images of spread spermatocytes from adult *Shoc1* KI homozygous (*Shoc1*^KI/KI^) mice and littermate controls (*Shoc1*^KI/+^) co-stained with SYCP3 (green) and REDIC1 (red). Scale bars, 10 μm. **(L)** Quantification of REDIC1 foci per cell at indicated meiotic stages using two-tailed Student’s t-test; **** *P* < 0.0001; error bars, mean ± SEM; n, the total number of nuclei analyzed. **(M)** Co-IP assays in HEK293T cells assessing interactions between human SHOC1-XPF (WT or mutant, FLAG-tagged) and M1AP/REDIC1/C1orf146 (HA/MYC-tagged). Input lysates were included. **(N)** Co-IP assays in HEK293T cells assessing interactions between mouse SHOC1-XPF (WT or mutant, FLAG-tagged) and M1AP/REDIC1/SPO16 (HA/MYC-tagged). Input lysates were included. **(O and P)** Relative interaction intensities quantified for human (O) or mouse (P) SHOC1-XPF (mutant) and M1AP / SHOC1-XPF (WT) and M1AP using two-tailed Student’s t-test; **** *P* < 0.0001; error bars, mean ± SEM. **(Q and R)** Relative interaction intensities quantified for human (Q) or mouse (R) SHOC1-XPF (mutant) and REDIC1 / SHOC1-XPF (WT) and REDIC1 using two-tailed Student’s t-test; ns, not significant; error bars, mean ± SEM. L-Zyg, late zygotene; Pach, pachytene.

Given the known DSBs repair defects in *Shoc1*^−/−^ mice(*14*), we then analyzed the expression of two essential recombinases, DMC1 and RAD51. In control spermatocytes, DMC1 and RAD51 localized to single-stranded DNA at the early-zygotene stage, and was removed from these sites during the late-zygotene to pachytene transition. At the early-pachytene stage, the number of DMC1 and RAD51 were reduced to approximately one-tenth compared to the early-zygotene stage. In *Shoc1* KI cells, DMC1 and RAD51 were similarly recruited to single-stranded DNA at the early-zygotene stages, and gradually slowly decreased from the early-zygotene stage to the pachytene stage. However, the number of both foci in pachytene KI spermatocytes were significantly higher than those in control spermatocytes (Figure S6A-D). Similarly, the number of SPATA22 and RPA2 foci in KI cells, which are well known for binding with single-stranded DNA overhangs during homologous recombination (HR), were also comparable to control ones in zygotene, but higher in pachytene spermatocytes (Figure S6E-H). Altogether, these results showed DSBs repair defects in *Shoc1* mutant cells.

During DSB recombination, ZMM proteins are recruited to recombination intermediates to facilitate the generation of COs-prone joint molecules, SEIs and double holiday junctions (dHJs). In control spermatocytes, TEX11 localized to recombination sites during meiotic prophase I, specifically marking the COs-prone recombination nodules in zygotene and early-pachytene spermatocytes. In contrast, TEX11 foci were absent in *Shoc1* KI cells (Figure 4E,F), indicating that the mutation within the XPF-like domain disrupted TEX11 localization. Additionally, MSH4, another component of the ZMM complex that facilitates the assurance and interference of COs, localized to mid-recombination intermediates in control early-pachytene meiocytes. Notably, a reduced number of MSH4 foci were observed in zygotene and pachytene spermatocytes of *Shoc1* KI males (Figure 4G,H), suggesting that the mutation within the XPF-like domain also influenced the recruitment of MSH4 foci to the chromosome axis and the stabilization of recombination intermediates. Intriguingly, the significantly decreased expression profiles in M1AP and REDIC1, two components of SHOC1 complex mentioned above, were also identified from late zygotene to pachytene stages in *Shoc1* KI males (Figure 4I-L). To define the interactions after the mutation within XPF-like domain, we generated FLAG-tagged SHOC1-XPF variants (p.Q646R) across species. Reciprocal co-IP assay was performed using anti-FLAG antibody, revealing a reciprocal interaction between M1AP, REDIC1, SPO16 and SHOC1 XPF-like domain in mice, whereas SHOC1 XPF-like domain could be interacted with M1AP, REDIC1 but not SPO16 in human. And the mutant SHOC1 (whether human or mouse origin) showed a significantly weaker interaction with M1AP instead of the other two proteins (Figure 4M-R). Altogether, the disruption of molecular interactions was associated with destabilization of the recombination intermediates in *Shoc1* KI mice.

The XPF-like domain in SHOC1 was reported to bind with branched DNA structures, which was essential for meiotic recombination. To elucidate the effect on interaction between branched recombination intermediates and the mutated XPF-like domain in SHOC1 (p.Q590R in human or p.Q646R in mice), we performed molecular dynamics simulations of WT and mutant SHOC1 proteins with the branched DNA (D-loop) structure (PDB ID: 7JY7). The human and mouse SHOC1/D-loop complexes were modeled using the HDOCK server with high confidence scores in WT (0.933 in human and 0.907 in mice) and KI groups (0.928 in human and 0.957 in mice) (Figure S7 A,B,F,G). Comparative analysis revealed a marked reduction in residue hydrogen bond formation mediated by human R590 (mouse R646) in the mutant complex relative to the WT configuration (Figure S7C,D,H,I). This substitution replaced the uncharged polar glutamine with a positively charged arginine, fundamentally altering local electrostatic properties that may perturb protein stability and intermolecular interactions. Quantitative analysis of docking scores derived from 100 top-ranked models per group further demonstrated significantly reduced binding energy for the mutant SHOC1/D-loop complex compared to WT (human: -190.6 ± 18.7 in p.Q590R vs. -202.2 ± 20.9 in WT, *P* < 0.0001; mouse: -189.4 ± 21.7 in p.Q646R vs. -203.5 ± 16.8 in WT, *P* < 0.0001) (Figure S7E,J). This substitution-induced conformational change appeared to mechanistically impair branched DNA binding ability of the XPF-like domain in SHOC1 during meiotic recombination. Thus, these data demonstrated that the XPF-like domain in SHOC1 ensures CO formation via orchestrating SHOC1 complex assembly and branched DNA structures binding.

### The disruption of dynamic 3D chromatin structure and CO region interaction in ***Shoc1* KI pachytene spermatocytes**

Chromatin undergoes dramatic 3D reorganization during meiosis (Figure 5A), most remarkably exemplified by the loop-axis reconfiguration of pachytene chromosomes. However, direct evidence linking chromatin architecture defects to meiotic arrest remains elusive. Thus, we isolated pachytene spermatocytes from *Shoc1* KI testes using a modified STA-PUT system (Figure S8A-C) and subsequently performed low-input Hi-C method (sisHi-C) to decipher chromatin architectural defects underlying meiotic arrest. The results of sisHi-C revealed significant alterations in the higher-order chromatin structure of *Shoc1* KI pachynema compared to the control group. Specifically, an overall decrease in distal interactions was observed in the mutant spermatocytes, as depicted in an interaction heatmap (Figure 5B). This observation was further supported by P(s) curve analysis, illustrating a reduced interaction frequency in distal regions, indicating distinct principles of chromatin folding (Figure 5C). In control pachynema, it was showed that distinct interchromosomal contact patterns was consistent with the alignment of chromosomes. Autosomal chromosomes in control pachytene stage still maintained strong A/B compartment identity observable in Hi-C contact map (Figure 5D left panel) and Pearson correlation matrices (Figure 5E left panel). Distinct from the autosomal pattern, the X chromosome’s compartment structure was significantly lost in pachynema. Consistent with the idea that the clustering interactions are linked to transcription, we observed a near-complete loss of this clustering on the X chromosome as it became transcriptionally silenced in pachynema (Figure 5F,G left panel). Previous data demonstrated that the pachytene chromatin organization is disrupted in *Sycp2*-mutant spermatocytes(*28*). Correspondingly, we observed increased compartmentalization of both autosome (Chromosome 2) and sex chromosome (Chromosome X) in *Shoc1* KI pachynema (Figure 5D-G right panel). In addition, the dynamics of topologically associated domains (TADs) were also analyzed. Compared to control spermatocytes, *Shoc1* KI pachynema exhibited more refined and segmented TADs, with increased numbers and a subtle enhancement of boundary insulation (Figure 5H-J).

**Figure 5.**
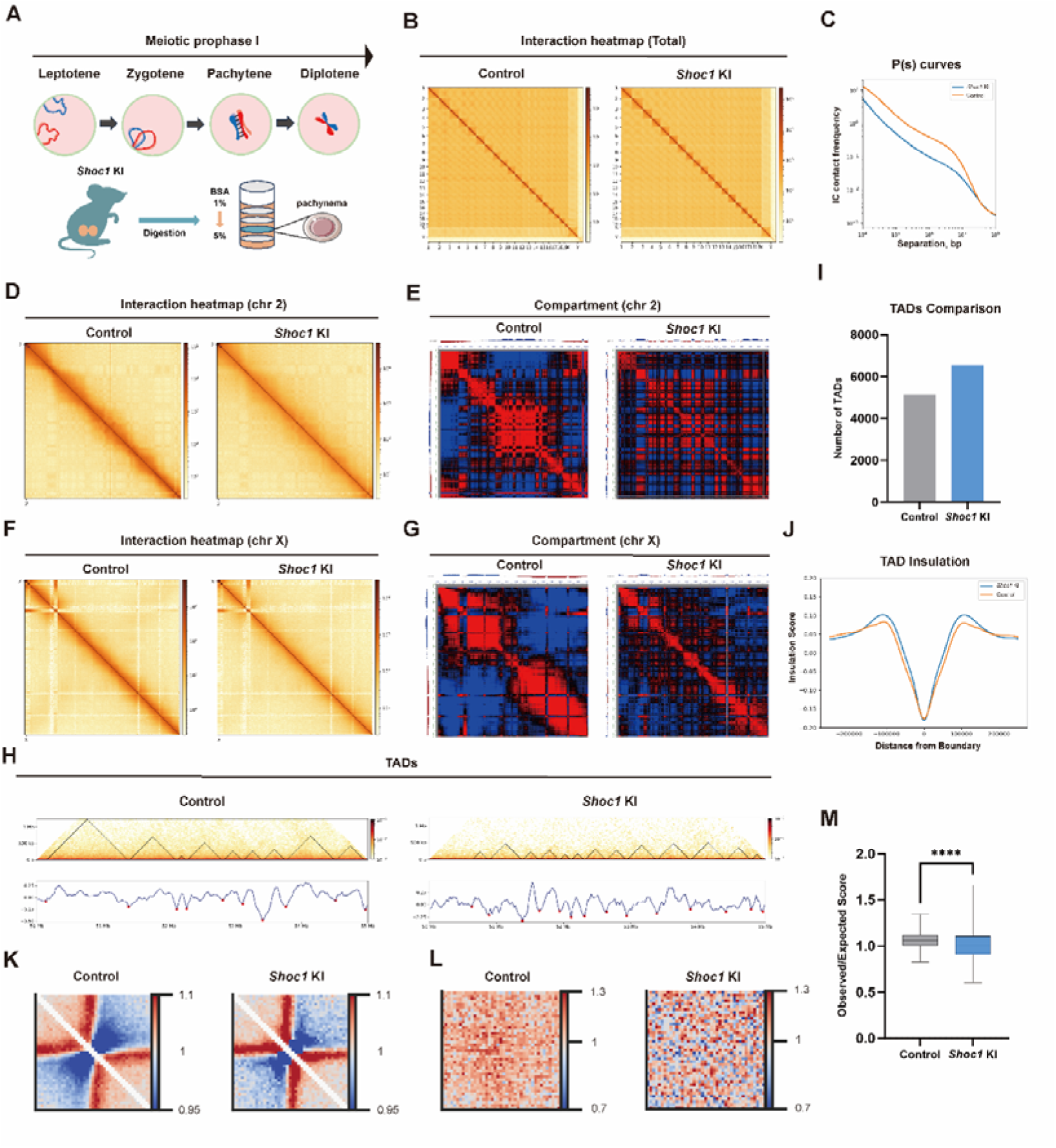
The disruption of dynamic 3D chromatin structure and CO region interaction in *Shoc1* KI pachytene spermatocytes. **(A)** Schematic of mouse meiotic prophase L and collection of pachytene spermatocytes by STA-PUT method. **(B)** Normalized Hi-C interaction heatmaps (2.5 MB bins) for all chromosomes in pachytene spermatocytes from adult *Shoc1* KI homozygous (*Shoc1*^KI/KI^) mice and controls (WT Hi-C dataset). **(C)** Chromatin contact probabilities relative to genomic distance (P(s) curves) between pachytene spermatocytes in *Shoc1* KI homozygous (*Shoc1*^KI/KI^) and control mice (WT Hi-C dataset). **(D)** Normalized Hi-C interaction heatmaps (500 kb bins) for chromosome 2 in pachytene spermatocytes from adult *Shoc1* KI homozygous (*Shoc1*^KI/KI^) mice and controls (WT Hi-C dataset). **(E)** Pearson’s correlation heatmaps (500 kb bins) showing A/B compartmentalization patterns for chromosome 2 in pachytene spermatocytes from adult *Shoc1* KI homozygous (*Shoc1*^KI/KI^) mice and controls (WT Hi-C dataset), with first principal component eigenvalues (PCA1) displayed. **(F)** Normalized Hi-C interaction heatmaps (500 kb bins) for chromosome X in pachytene spermatocytes from adult *Shoc1* KI homozygous (*Shoc1*^KI/KI^) mice and controls (WT Hi-C dataset). **(G)** Pearson’s correlation heatmaps (500 kb bins) showing A/B compartmentalization patterns for chromosome X in pachytene spermatocytes from adult *Shoc1* KI homozygous (*Shoc1*^KI/KI^) mice and controls (WT Hi-C dataset), with PCA1 displayed. **(H)** Hi-C interaction matrix of chromosome 1 (50-55 Mb; 25 kb resolution) in pachytene spermatocytes from adult *Shoc1* KI homozygous (*Shoc1*^KI/KI^) mice and controls (WT Hi-C dataset). Triangles denote TADs. **(I)** TADs count in pachytene spermatocytes from adult *Shoc1* KI homozygous (*Shoc1*^KI/KI^) mice and controls (WT Hi-C dataset). **(J)** Average insulation scores ±250 kb from TADs boundaries (25 kb resolution) in pachytene spermatocytes from adult *Shoc1* KI homozygous (*Shoc1*^KI/KI^) mice and controls (WT Hi-C dataset). **(K and L)** Aggregated Hi-C interaction heatmaps flanking TAD boundaries (500 kb regions; K) or centered at CO-associated DSBs (2 Mb regions; L) in pachytene spermatocytes from adult *Shoc1* KI homozygous (*Shoc1*^KI/KI^) mice and controls (WT Hi-C dataset). Interactions normalized as observed/expected (Obs/Exp; 5 kb bins). **(L)** Box plots comparing Obs/Exp Hi-C interactions in 2 Mb regions centered at CO-associated DSBs in pachytene spermatocytes from adult *Shoc1* KI homozygous (*Shoc1*^KI/KI^) mice and controls (WT Hi-C dataset) using two-tailed Student’s t-test; **** *P* < 0.0001; error bars, mean ± SEM.

During early meiotic prophase, DSBs emerged preferentially at chromatin loops anchorage sites along the chromosome axis, thereby facilitating homologous chromosome co-alignment. Notably, genomic regions with DSBs, particularly those linked with CO-designated DSBs (CO-DSBs), exhibited substantial reorganization of chromatin interaction networks concurrent with homologous pairing and axial alignment processes(*29*). To investigate the spatiotemporal relationship between CO formation and chromatin architecture, we mapped CO hotspots using established datasets combined with high-resolution sisHi-C interaction analysis in *Shoc1* KI spermatocytes. Pileup heatmap visualization revealed a significant reduction in chromatin interaction frequency flanking CO sites in the pachytene-stage nuclei of *Shoc1* KI mice compared to control regions (Figure 5K-M). This attenuation of local chromatin connectivity near CO hotspots mirrored the dynamic chromatin reorganization patterns typically observed in zygotene spermatocytes(*29-32*), suggesting that *Shoc1* mutation perturbs normal chromatin compaction dynamics during pachytene progression. These findings demonstrated that the variant within the XPF-like domain in SHOC impaired dynamic 3D chromatin structure and CO region interaction.

### *Shoc1* KI spermatocytes exhibited incomplete synapsis

Since synapsis and HR are interdependent and closely coupled processes during meiotic prophase, we subsequently investigated the dynamic of chromosomal synapsis in *Shoc1* KI spermatocytes. We first stained the spermatocyte spreads with antibodies against SYCP3 and HORMAD1, which is localized along the unsynapsed chromosome axis in pachynema. In control mice, the HORMAD1 signal was observed only at unsynapsed sex chromosomal regions in the pachytene stage. However, in *Shoc1* KI spermatocytes, persistent HORMAD1 signals were identified on the autosomal chromosome axis (Figure 6A). After further staining the spermatocyte spreads for SYCP3 and SYCP1 (a transverse filament of the SC), as expected, the SYCP1 signals extended to the entire length of the autosomal axes and the pseudoautosomal region (PAR) of the sex chromosomes in control pachytene spermatocytes. In *Shoc1* KI cells, the majority of pachytene-like cells with short and thick lateral elements displayed discontinuous and/or weak SYCP1 signals, suggesting that synaptic defects occur after the mutation within XPF-like domain in *Shoc1* (Figure 6B,D).

**Figure 6.**
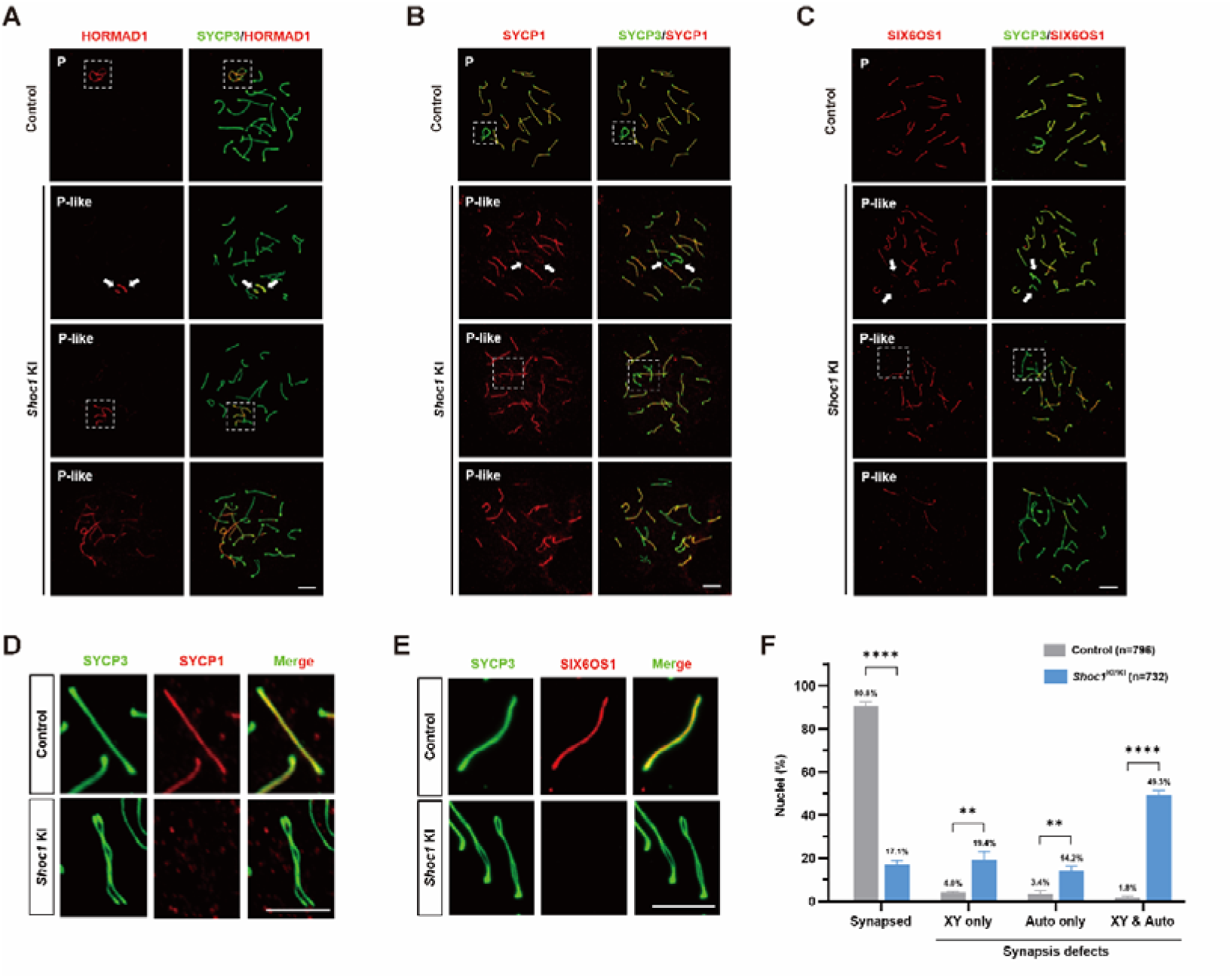
*Shoc1* KI spermatocytes exhibited incomplete synapsis. **(A-C)** Representative images of spread spermatocytes from adult *Shoc1* KI homozygous (*Shoc1*^KI/KI^) mice and littermate controls (*Shoc1*^KI/+^) co-stained with SYCP3 (green) and (A) HORMAD1 (red), (B) SYCP1 (red), (C) SIX6OS1 (red). Scale bars, 10 μm. **(D and E)** Enlarged views of spread spermatocytes co-stained with SYCP3 (green) and (D) SYCP1 (red), (E) SIX6OS1 (red). Scale bars, 10 μm. **(F)** Quantification of synaptic defects in pachytene spermatocytes from adult *Shoc1* KI homozygous (*Shoc1*^KI/KI^) mice and littermate controls (*Shoc1*^KI/+^) using two-tailed Student’s t-test; **** *P* < 0.0001; ** *P* < 0.01; error bars, mean ± SEM; n, the total number of nuclei analyzed. XY, X and Y chromosomes; Auto, autosomes.

Further, we stained the spermatocyte spreads with antibodies against SYCP3 and the SC central element protein SIX6OS1 (Figure 6C,E). Similar to the observation in SYCP1 staining spermatocytes, discontinuous SIX6OS1 signals were evident in the pachytene spermatocytes of *Shoc1* KI mice. Remarkably, in control mice, 90.8 ± 1.5% of pachytene spermatocytes achieved complete synapsis. However, in *Shoc1* KI mice, although 17.1 ± 1.6% of pachytene spermatocytes presented fully synapsed chromosomes, the remaining showed variable synaptic abnormalities. Among them, 14.2 ± 1.7% of cells showed synapsed sex chromosomes but with at least one pair of incompletely synapsed autosomes, 19.4 ± 3.0% of cells showed fully synapsed autosomes but with asynapsed sex chromosomes, and 49.3 ± 1.6% contained synapsis defects on both autosomal and sex chromosomes (Figure 6F). Thus, these findings indicated that *Shoc1* mutation (p.Q646R) could lead to synaptic defects.

### The XPF-like domain in SHOC1 establishes a molecular barrier safeguarding autosome from meiotic silencing of unsynapsed chromatin (MSUC)

During meiosis, unsynapsed chromosomal regions are transcriptionally silenced through a conserved mechanism known as MSUC. Specifically, the silencing of unsynapsed sex chromosomes is termed meiotic sex chromosome inactivation (MSCI)(*33, 34*). Varying degrees of asynapsis could result in disturbance of the normal loading of MSUC proteins, impacting gene expression on autosomal and sex chromosomes, leading to extensive spermatocyte apoptosis(*35, 36*). To investigate whether the MSUC is disrupted in *Shoc1* KI pachytene-like spermatocytes, RNA polymerase II (POL II) was used to detect transcription activity within chromosomes. In control pachynema, POL II signals were distinctly enriched on autosomes consistent with transcriptional silencing via MSCI. In contrast, *Shoc1* KI cells exhibited a pronounced loss of POL II signals not only on sex chromosomes but also in adjacent autosomal regions (39.0 ± 1.7% in KI vs. 3.1 ± 0.6% in control, *P* < 0.0001) (Figure S9A,B). Since the DDR pathway is essential for establishing and maintaining MSCI, we measured MDC1, a DDR factor critical for the amplification of DDR factors from the axis to the chromosome-wide domain(*33*), in *Shoc1* KI mice. Correspondingly, a higher ratio of trapped MDC1 signals were observed on both sexual chromosomes and autosomes in the KI group compared to the control group (53.0 ± 3.0% in KI vs. 1.6 ± 0.5% in control, *P* < 0.0001) (Figure S9C,D). These results suggested that the overloaded MSUC in *Shoc1* KI pachytene-like spermatocytes could impede the expression of genes essential for meiotic progression, potentially contributing to the MML arrest.

To confirm the association of the XPF-like domain in SHOC1 with MSUC and determine the consequences of overloaded MSUC in *Shoc1* KI mice, we compared the transcriptome of KI pachynema with control cells using 10 × Genomics scRNA-Seq. Distinct germ cell types were identified based on Uniform Manifold Approximation and Projection (UMAP) and marker gene analysis from both control and *Shoc1* KI testes (Figure 7A-B). The cell number as well as proportion for each cell cluster were summarized (Figure 7C). Notably, the findings revealed that the KI group lacked post-meiotic haploid spermatids, and the proportion of various substages of spermatocytes during meiosis was similar to our results from spread staining. With the cutoff of log2(fold change) ≥0.25 or ≤ −0.25, a total of 645 differentially expressed genes (DEGs), including 527 down-regulated genes and 118 up-regulated genes in *Shoc1* KI pachytene-like spermatocytes compared with WT, were identified (Figure 7D). Gene Set Enrichment Analysis (GSEA) of downregulated genes in the *Shoc1* KI compared with control pachytene spermatocytes include “nucleus organization”, “germ cell development”, “cilium or flagellum-dependent cell motility”, “chromosome condensation” and “spermatid differentiation” (Figure 7E). Given that the *Shoc1* KI pachytene spermatocytes exhibited overloaded MSUC, we compared mRNA levels between autosomes, X chromosome, and Y chromosome was performed. We found that the majority of DEGs were distributed on the autosomes (619/645, 96.0%) rather than the sex chromosomes (X and Y chromosomes) (Figure 7F). Intriguingly, genomic transcription mapping across chromosomes revealed that autosomal downregulated genes were predominantly clustered on chromosome 16. Functional annotation demonstrated strong association of these chr16 genes with critical reproductive processes including “chromosome condensation”, “spermatid differentiation”, and “germ cell development” through rigorous GO enrichment analyses (Figure 7G,H), mechanistically linking this dysregulation to apoptosis during MMI and meiotic arrest. Unexpectedly, a lower level of gene expression on sex chromosomes was also observed in *Shoc1* KI pachytene spermatocytes (Figure 7I), suggesting that SHOC1 may play a role in regulating MSCI as well. Overall, our findings revealed that the XPF-like domain in SHOC1 establishes a molecular barrier preventing autosome intrusion into the sex body compartment, thereby safeguarding critical autosomal loci from MSUC. Additionally, chromosome 16 exhibits heightened MSUC vulnerability, as shown by its disproportionate transcriptional suppression in *Shoc1* mutation within XPF-like domain.

**Figure 7.**
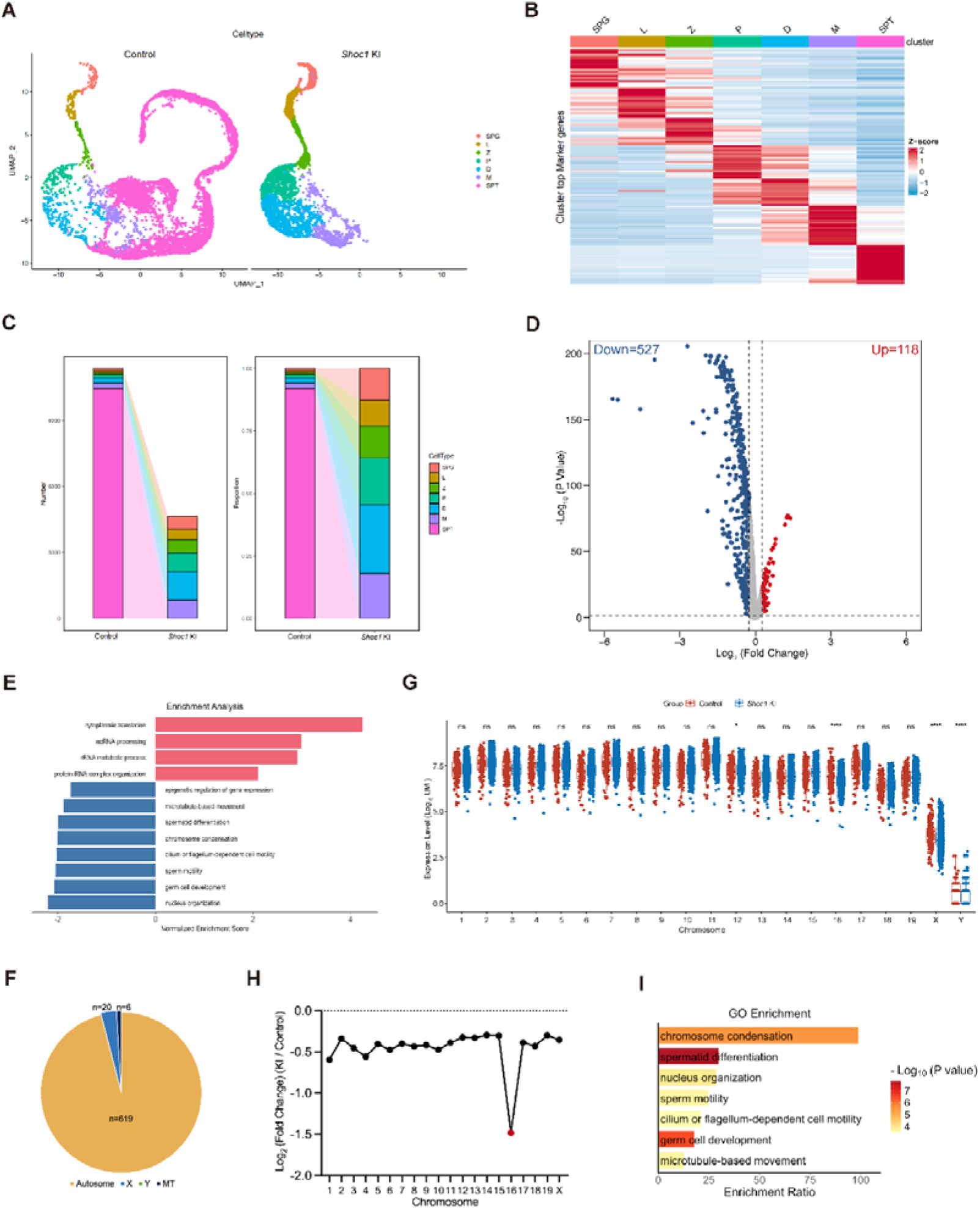
*Shoc1* KI led to overloaded MSUC and autosome-linked gene silencing in pachytene spermatocytes. **(A)** UMAP plots of testicular single-cell transcriptomes from adult *Shoc1* KI homozygous (*Shoc1*^KI/KI^) mice and littermate controls (*Shoc1*^KI/+^). **(B)** Heatmap displaying the top DEGs across germ cell subclusters. A gradient of blue to red indicates low to high expression levels. **(C)** Stacked bar charts quantifying cell type proportions and counts from adult *Shoc1* KI homozygous (*Shoc1*^KI/KI^) mice and littermate controls (*Shoc1*^KI/+^). SPG: spermatogonia; L: leptotene spermatocyte; Z: zygotene spermatocyte; P: pachytene spermatocyte; D: diplotene spermatocyte; M: metaphase spermatocyte; SPT: spermatid. **(D)** Volcano plot of DEGs in pachytene spermatocytes from adult *Shoc1* KI homozygous (*Shoc1*^KI/KI^) mice and littermate controls (*Shoc1*^KI/+^); Cut-off: log2(Fold Change) ≥ 0.25 or ≤ −0.25, *P* < 0.05. **(E)** GSEA of downregulated (blue) and upregulated (red) DEGs in pachytene spermatocytes from adult *Shoc1* KI homozygous (*Shoc1*^KI/KI^) mice and littermate controls (*Shoc1*^KI/+^). **(F)** Distribution of DEGs across autosomes, sex chromosome (X), and mitochondrial genome (MT). **(G)** Normalized UMI counts for genes on each chromosome across spermatogenesis stages using two-tailed Student’s t-test; * *P* < 0.05; **** *P* < 0.0001; ns, not significant; error bars, mean ± SEM. **(H)** Mean log2(fold change) of autosomal and X chromosome gene expression. **(I)** GO analysis of downregulated DEGs on chromosome 16 in pachytene spermatocytes from adult *Shoc1* KI homozygous (*Shoc1*^KI/KI^) mice and littermate controls (*Shoc1*^KI/+^).

**Figure 8.**
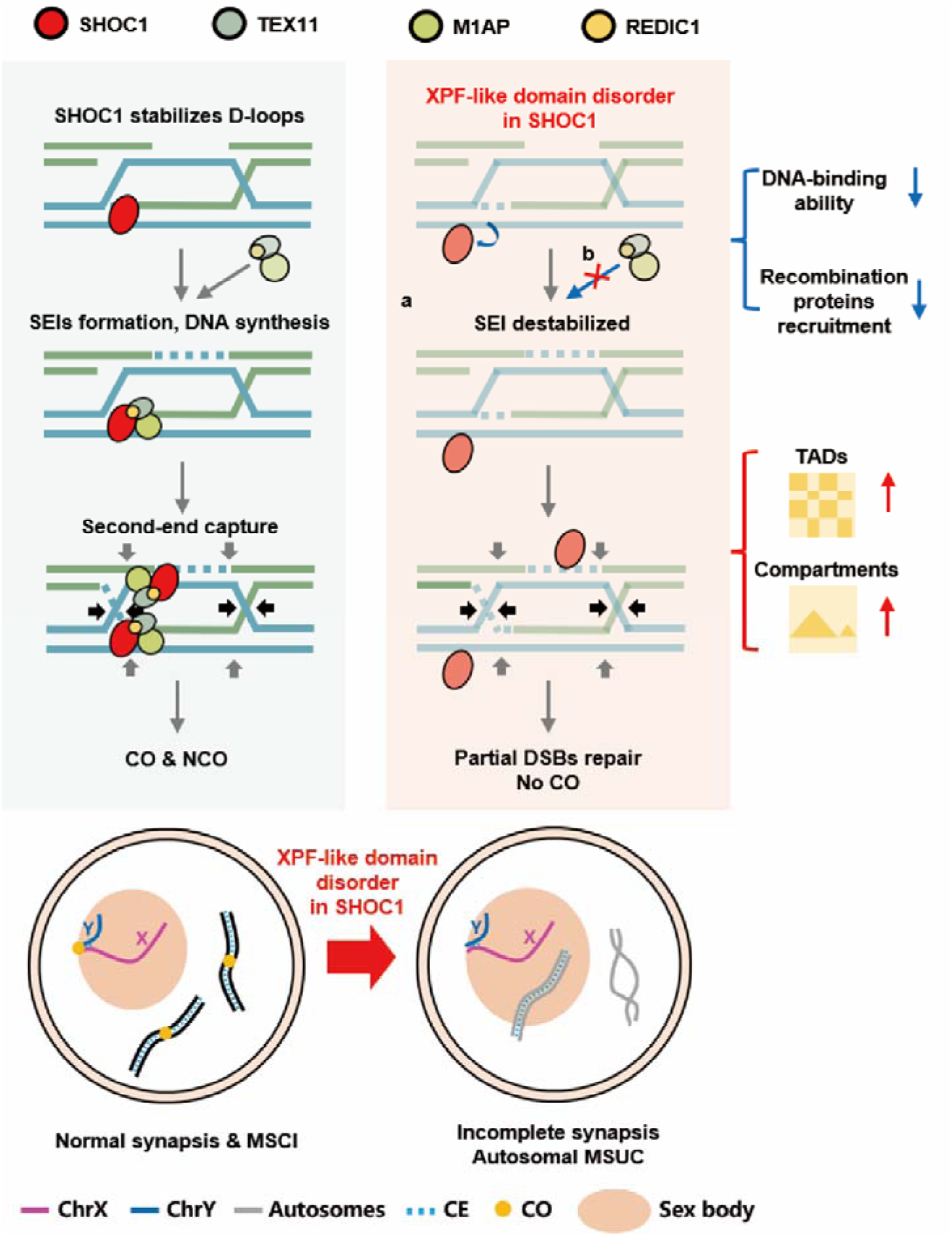
Illustration of the XPF-like domain’s role in SHOC1-mediated homologous recombination and safeguarding autosome from MSUC in meiosis. This study revealed the role of the XPF-like domain in SHOC1 as a molecular scaffold regulating synapsis and CO formation during meiosis. The p.Q590R variant within the XPF-like domain of SHOC1 impaired DSBs repair by compromising the SHOC1’s ability to bind branched DNA structures and the recruitment of M1AP, REDIC1, and ZMM factors to recombination intermediates, ultimately abolishing CO formation and causing meiotic arrest. Hi-C analyses revealed that mutant pachytene spermatocytes exhibited compromised chromatin architecture – increased TADs insulation and elevated compartmentalization. Furthermore, the XPF-like domain in SHOC1 was indispensable for safeguarding autosome from MSUC in pachytene spermatocytes.

## DISCUSSION

As the foundation of sexual reproduction, meiosis is required to ensure genome stability and heritable diversity by generating haploid gametes through the formation of DSBs, homologous pairing, synapsis, HR, and CO formation. Defects in evolutionarily molecular machines can result in meiotic arrest or aneuploidy, ultimately damaging human fertility. Our findings revealed the critical role of the XPF-like domain in SHOC1 as a molecular scaffold regulating CO formation and synapsis during meiosis. Species-specific interactome analyses identified distinct SHOC1 complexes between human and mice. The p.Q590R variant within the XPF-like domain in *SHOC1* impaired DSBs repair by compromising the SHOC1’s ability to bind branched DNA structures and the recruitment of M1AP, REDIC1, as well as ZMM factors to recombination intermediates, ultimately abolishing CO formation and causing meiotic arrest. Notably, Hi-C analyses revealed that mutant pachytene spermatocytes exhibit disrupted chromatin architecture–increased TADs insulation, elevated compartmentalization, and weakened interactions at CO sites–phenocopying zygotene-like chromatin states. We further identified SHOC1’s novel role in safeguarding autosomes from the XY body, with transcriptomic data showing chromosome 16-specific silencing of meiotic regulators through overloaded MSUC activation.

In the conserved Zip2-Zip4-Spo16 (ZZS) trimeric complex of yeast, Zip4 interacts with the N-terminal domain of ZIP2, while SPO16 binds specifically to its C-terminal XPF-like domain, which is required for the formation of class L interfering COs and polymerization of the SC(*37, 38*). Recent studies reported two novel proteins M1AP and REDIC1 in mouse testes can interact and cooperate with the ZZS complex in the stabilization of recombination intermediates(*22, 23*). To further confirm the interaction between SHOC1 and these two proteins and other ZZS components, we performed co-IP in lysates of HEK293T cells co-expressing human or mouse SHOC1with TEX11, SPO16, M1AP or REDIC1. The results uncovered previously unrecognized species-specific interactome in SHOC1 complex that redefined its function in human meiosis. The absence of direct SHOC1-C1orf146 interaction and the absolute dependence on an intact N-terminal domain for TEX11 binding in human revealed a species-specific structural scaffold deciphering regulation of human CO formation. Further complexity of SHOC1’s functional network emerges from the integration of M1AP/REDIC1, two novel interactors absent in canonical ZMM models. These findings transcend the classical “ZMM stabilizer” paradigm, positioning SHOC1 complex as a central integrator of DSBs repair and chromosome-scale dynamics.

Several reports have identified the clinical relevance of *SHOC1* variants in varying severity of meiotic arrest phenotypes. Wang *et al.* reported their findings of bi-allelic *SHOC1* variants in three NOA-affected patients, of which two are homozygous for the same LoF variant (c.231_232del:p.L78Sfs*9), and one is heterozygous for two different missense variants (c.1978G>A:p.A660T; c.4274G>A:p.R1425H). Testicular biopsy of one patient revealed impairment of spermatocyte maturation(*26*). In contrast, Csilla *et al.* documented 4.2% of spermatocytes in a patient homozygous for a frameshift variant c.797delT (p.L266Qfs*6) achieving XY body formation and progressing to MMI*(39)*. Here, we expanded the genetic landscape of meiotic arrest by screening for pathogenic *SHOC1* variants and revealed severe zygotene arrest without XY body formation in the testicular section of probands. The abnormalities were similar to those in germ-cell specific (conditional knockout) cKO and systemic KO mouse models arrested at zygotene-like stage(*14, 26*). For several of the identified variants, we demonstrated functional data linking the impaired integrity of encoded proteins due to different frameshift types or aberrant mRNA splicing to downstream cellular effects. Importantly, our results provided strong evidence for the clinical relevance and emphasized the necessity for *SHOC1* gene screening in NOA diagnostics.

Given the high evolutionary conservation of the XPF-like domain in SHOC1 across species, investigation toward the critical residues and systematic characterization of molecular mechanisms becomes imperative to elucidate the XPF-like domain’s essential roles during meiotic progression. While previous investigations have primarily utilized bioinformatic analyses and *in vitro* biochemical approaches(*17, 37*), direct *in vivo* evidence remains limited. In this study, we identified a missense variant within the XPF-like domain (p.Q590R) through genetic screening and established a CRISPR/Cas9-edited KI mouse model, which exhibited not only COs abolishment but also synaptic defects. Recent work on a complete SHOC1-null mouse model revealed even more severe phenotypes, showing complete arrest at a zygotene-like stage with defective synapsis, unresolved DSBs, and absence of both ZMM protein-associated recombination intermediates and MLH1-marked class L COs(*14, 16*). Comparatively, our KI model and the reported *Shoc1*^hyp/hyp^ mutant (expressing reduced levels of truncated SHOC1) (*17*)both displayed MMI arrest with univalent chromosomes. This phenotype closely resembles those observed in TEX11(*40*), M1AP(*22*), and REDIC1(*23*) deficient mouse models, collectively suggesting that the functional role of the Q646 residue (corresponding to human Q590 residue) within the XPF-like domain in SHOC1 aligns with the critical transition from early recombination intermediates to mature COs during late prophase I.

The more severe synapsis defects observed in our *Shoc1* KI mice compared to these other models underscored the central role of SHOC1 complex as a molecular scaffold in regulation both synapsis and CO formation during meiosis. It has been reported that mutations in multiple DNA-binding surfaces on Zip2-Spo16 severely compromise DNA binding, supporting a model in which the XPF-like domain cooperates to bind with branched DNA structures(*37*). Thus, we investigated the effects resulting from the current mutation within the XPF-like domain on its ability to bind branched DNA structures using molecular docking. The findings demonstrated that the substitution-induced conformational changes appeared to mechanistically impair branched DNA (D-loop) binding ability of the XPF-like domain in both human and mouse SHOC1.

Chromatin undergoes drastic reprogramming during meiosis(*29-32*). Previous studies have examined dynamics of the 3D meiotic chromatin architecture in mice and rhesus monkey, in which chromosomal TADs undergo dissolution and re-establishment during spermatogenesis(*28, 31*). This is accompanied by the emergence of a unique chromatin configuration as the SC forms, featured by highly refined, transcriptionally correlated chromatin compartments. Notably, the fact that both SYCP2 and TOP6BL deficient mice, where the SC failed to be established, exhibited the restoration of TADs and increased conventional compartmentalization in spermatocytes, suggesting that the SC may restrict TADs and promote local compartments(*28*). In the present study, sisHi-C analyses unveiled that *Shoc1* KI disrupts hierarchical chromatin organization, concomitant with meiotic arrest. While compartment/TADs features persist in *Shoc1* mutant pachynema, diminished distal interactions and failed chromosome alignment revealed SHOC1’s essential function in coordinating higher-order chromatin topology. Moreover, the attenuation of chromatin connectivity at CO hotspots in KI group suggested that SHOC1 could safeguard local chromatin compaction during pachytene, ensuring proper coordination between SC assembly and homology-directed DSBs repair.

Intriguingly, our investigation showed that overloaded MSUC took place in pachytene spermatocytes of *Shoc1* KI mice. Combined with scRNA sequencing analysis, the KI group exhibited autosomal transcriptional suppression, with chromosome 16 displaying the most significant differential expression relative to control counterparts. Significantly, chromosome 16 contains numerous critical genes in regulating spermatogenesis, potentially explaining its preferential transcriptional vulnerability during meiotic arrest. This chromosomal predisposition holds clinical relevance, as trisomy 16 represents a frequent autosomal anomaly in early gestational loss(*41*). While these data suggested chromosomal specificity in transcriptional perturbation, the precise mechanisms underlying how the XPF-like domain mutation in *SHOC1* mediates autosomal MSUC overload remain to be fully elucidated.

In conclusion, our findings delineated species-specific divergences between human and mouse SHOC1 complex, and revealed the critical role of the XPF-like domain in SHOC1 as a molecular scaffold regulating synapsis and CO formation during meiosis. The disorder of the XPF-like domain in SHOC1 impaired DSBs repair by compromising its ability to bind branched DNA structures and the recruitment of M1AP, REDIC1, and ZMM factors to recombination intermediates, ultimately abolishing CO formation. Furthermore, it disrupted dynamic 3D chromatin structure in pachytene spermatocytes and induced defects in homologous chromosome synapsis. More importantly, the XPF-like domain in SHOC1 was revealed to prevent autosome intrusion into the sex body compartment, thereby safeguarding critical autosomal loci from MSUC.

## MATERIALS AND METHODS

### Study participants

The experiments performed in human were approved by the ethics committee of Shanghai General Hospital (2022SQ294). Written informed consent was obtained from the donors for the use of their clinical data and testicular tissues for research purposes. For the initial cohort, a total of 1072 idiopathic NOA (iNOA) patients were enrolled from the Department of Andrology, Urologic Medical Center, Shanghai General Hospital, Shanghai Jiao Tong University School of Medicine, Shanghai, China. Physical examination and repeated semen analyses following centrifugation were conducted and assessed in accordance with the guidelines of the World Health Organization guidelines (5^th^ edition)(*42*). Known causal factors for male infertility, including orchitis, cryptorchidism, varicocele, radiation, chemotherapy, and testicular cancer were excluded. The genetic screening, including karyotype and Y chromosome microdeletions, was performed in the patients with NOA. The definition of the testicular phenotype was based on multiple testis biopsies (one fragment was analyzed by the pathologist who described the histological picture, the remaining fragments were analyzed by the embryologist who searched for spermatozoa). Ultimately, 171 patients with meiotic arrest were subjected to whole-exome sequencing (WES) in this study. Additionally, patients with obstructive azoospermia (OA), characterized by normal spermatogenesis but seminal tract obstruction, were included as a positive control.

### WES and Sanger sequencing

Genomic DNA was isolated from peripheral blood samples using a TIANamp Blood DNA Kit (TIANGEN) according to the manufacturer’s protocol. gDNA was subjected to WES performed by the company (Shanghai Yuyin Biotechnology Co., Ltd) on the HiSeq2000 sequencing platform (Illumina Inc., San Diego, CA, USA), as described previously(*43, 44*). WES data analysis was performed using the Genome Analysis Toolkit. Briefly, the WES raw reads after removing adaptors were aligned to the human genome (GRCh37/hg19) using the Burrows-Wheeler Aligner(*45*), followed by removal of the PCR duplicates and sorting using Picard (http://broadinstitute.github.io/picard/). Variant identification was performed using the Genome Analysis Toolkit package following the recommended best practices, including base recalibration variant calling with Haplotype Caller, variant quality score recalibration and variant annotation using the ANNOVAR software. Candidate causative variants that matched the following criteria were selected: (i) frequency below 1% in public human databases (1000 Genomes mutation database and gnomAD); (ii) predicted to be deleterious variants according to multiple bioinformatics tools (SIFT, PolyPhen-2 and MutationTaster); (iii) homozygous variants were prioritized; and (iv) relevancy for infertile phenotype using comprehensive expression data (testis-enriched) (http://www.proteinatlas.org/) and model organism data (http://www.informatics.jax.org/mgihome/homepages/).

The *SHOC1* variants identified by WES were further validated by Sanger sequencing. Polymerase chain reaction (PCR) was conducted, and the primers are listed in supplementary table S1. Subsequently, the PCR products underwent bidirectional Sanger sequencing using a 3730xl DNA Analyzer (Applied Biosystems, Foster City, California, USA) in Tsingke Life Technologies Biotechnology Co., Ltd. (Tsingke, Beijing, China).

### Mice

The animal experiments were approved by the Experimental Animal Management and Ethics Committee (2022AWS0287), and complied with the guidelines of the Animal Care and Use of Shanghai General Hospital, Shanghai Jiao Tong University School of Medicine. *Shoc1* KI mice, corresponding to the *SHOC1* variant (c.A1769G) identified in the patient (P21226), were generated using CRISPR/Cas9 genome editing technology. Briefly, Cas9 mRNA, single-guide RNAs (sgRNAs), and oligonucleotides were co-injected into the zygotes of C57BL/6 mice, followed by embryo transfer into pseudopregnant ICR females. Genomic DNA was extracted from the tails of founder mice, and genotyping was performed by PCR and Sanger sequencing. The founder mice carrying the target point mutation were crossed with WT C57BL/6 mice to obtain offspring. PCR assays and Sanger sequencing were used to identify the point mutation in founder mice. The sequences of sgRNAs, donor oligo and the primers for genotyping are shown in supplementary table S1.

### Histological analysis and immunofluorescence (IF) staining

Testes, epididymis and ovaries were detached and immediately fixed overnight in 4% paraformaldehyde. For histological analysis, sections were processed and stained with H&E according to standard protocols as described previously(*26*). For immunostaining of the sections, the tissue sections were dewaxed in xylene, re-hydrated in a descending alcohol gradient, and heated in sodium citrate buffer (90-98°C) for 15 min for antigen retrieval. After blocking with 5% normal donkey serum (NDS, 017-000-121, Jackson) for 1 h at room temperature (RT), the sections were incubated overnight with primary antibodies at 4°C. The sections were washed three times with phosphate-buffered saline containing Tween-20 (PBST), and incubated with highly cross-adsorbed secondary antibodies conjugated with Alexa Fluor^®^ 488 or Alexa Fluor^®^ 594 for 1 h at RT. The sections were washed three times with PBST and counterstained with DAPI (H-1200, Vector) to label the nuclei. Sections of the samples were imaged on an FV3000 confocal microscope (Olympus). The details of primary and secondary antibodies are listed in supplementary table S2.

### Plasmid construction and mutagenesis

Total RNA was extracted from testes, HEK293T and HeLa cells using TRIzol reagent (15596026, Thermo Fisher Scientific) according to the manufacturer’s procedure and were reverse-transcribed to generate cDNA with SuperScript IV Reverse Transcriptase (6215A, TaKaRa). The cDNAs encoding all included proteins were cloned by PCR amplification from the human or mouse testis cDNA library with the PrimeSTAR system (R045A, TaKaRa). Sequences were inserted into plasmids via HR using a ClonExpress MultiS One Step Cloning Kit (C113, Vazyme). All sequences cloned into vectors were fully sequenced and then analyzed using BLAST to confirm the correct insertion of the sequences. All fusion proteins were designed to prevent the generation of frameshift-mutant proteins. Primers for constructing plasmids in this paper are listed in supplementary table S1. In addition, the truncated SHOC1 plasmids (human and mouse SHOC1 ΔF1-ΔF4) were generated by Tsingke Life Technologies Biotechnology Co., Ltd. (Tsingke, Beijing, China).

### Cell culture and transfection

HEK293T or HeLa cells were cultured in DMEM/high glucose (SH30243.FS, Cytiva) supplemented with 10% FBS (10099141C, Gibco) and 1% penicillin-streptomycin (15140122, Gibco) at 37L in 5% CO_2_. For each well of a 6-well plate, 2.5 μg of plasmids and 5 μl of Lipofectamine 3000 (L3000015, Invitrogen) were used for transfection. Cells were collected 48 h after transfection for further analysis.

### Minigene assay

The variant of *SHOC1* (c.2738-1G>A) is located at the splice-site acceptor of intron 21. We acquired the [Intron (179 bp)-Exon 21 (118 bp)-Intron (112 bp)] region by PCR and cloned it into a modified pcMINI plasmid. The variant (c.2738-1G>A) plasmid was generated by PCR with WT plasmid as the template. The WT and mutant plasmids were transfected into HEK293T cells. After culture for 48 h, total RNA was extracted using TRIzol and reverse-transcribed to obtain cDNA. RT-PCR products were separated by electrophoresis on 2% agarose gels containing ethidium bromide and visualized by exposure to UV light. Each DNA band was gel-extracted with the use of Gel Extraction Kit (D2500, Omega) and sequenced.

### Modeling of SHOC1/D-loop complex structure through molecular docking

The SHOC1/D-loop complexes were modeled using the HDOCK server(*46*). Structural model prediction of WT and mutant (p.Q590R in human and p.Q646R in mice) SHOC1 protein were anticipated in AlphaFold3 (https://golgi.sandbox.google.com/). The structure of D-loop was obtained from a previously solved structure of RecA-D-loop complexes (PDB ID: 7JY7)(*47*) and used for this docking study. The default docking parameters provided by HDOCK server were used and the docking conformations derived from 100 top-ranked models per group were further analyzed. All structural of figures were done using UCSF Chimera (https://www.rbvi.ucsf.edu/chimerax).

### Co-immunoprecipitation (co-IP)

The testicular tissue or cells were lysed in IP lysis buffer (20 mM HEPES, 150 mM NaCl, 0.5% NP-40, 1 mM DTT, pH = 7.3) with 1 × protease inhibitor cocktail (11836153001, Roche) for 30 min on ice. The lysates were clarified by centrifugation at 12,000 × g for 20 min at 4 °C. Next, 10% of the supernatants were kept to be used as the input control, and the remainder was incubated with relevant antibodies overnight on a shaker. The lysates (500-1000 μg) were then incubated with either FLAG-beads (F2426, Sigma-Aldrich) or protein A/G magnetic beads (88803, Life Technologies) under rotation at 4°C for 2 h. The proteins associated with the protein A/G magnetic beads were collected by placing the tubes into a magnetic stand and rinsed five times with immunoprecipitation washing buLer (10 mM Tris–HCl pH 7.5, 1 mM EDTA, 150 mM NaCl, and 0.1% Triton X-100), after which they were subjected to subsequent operation.

### Western blot (WB)

The testicular tissue or cells were rinsed with PBS and lysed in RIPA lysis buffer (89901, Sigma-Aldrich) with 1 × protease inhibitor cocktail (11836153001, Roche) for 30 min on ice. Following centrifugation at 12,000 g for 20 min at 4°C, the protein concentration of the lysates was determined using a BCA kit (23225, Thermo Fisher Scientific). SDS-PAGE (Sodium dodecyl sulfate-polyacrylamide gel electrophoresis) was performed using 20 μg of lysate from each sample, and WB was carried out as previously described(*48*). After conducting SDS-PAGE on 10% gel at 80 V for 30 min followed by 120 V for 80 min to load and separate the protein samples, the samples were transferred to a 0.22 μm PVDF membrane (ISEQ00010, Millipore). The membranes were then blocked with 5% skim milk and incubated with the primary antibodies overnight at 4°C. The next day, the membranes were washed three times with TBS containing 0.1% Tween (TBST) and then incubated with horseradish peroxidase-conjugated secondary antibodies for 1 h at RT. After three washes in TBST, the blots were detected by chemiluminescence (WBKLS0500, Millipore). The densitometry was analyzed using the AI600 software. The housekeeping protein β-actin was used for WB normalization. The details of primary and secondary antibodies are listed in supplementary table S2.

### Spermatocyte spreads and immunostaining

Mouse testicular cells were prepared for surface spreading and subsequent immunostaining as previously described with the following modifications. The testicular tissue was macerated in PBS, followed by removal of the tunica albuginea. The seminiferous tubules were incubated in hypotonic extraction buffer (0.6 M Tris pH 8.2, 500 μl; 0.5 M sucrose, 1 ml; 0.17 M trisodium citrate dihydrate, 1 ml; 0.5 M EDTA pH 8.0, 100 μl; 0.5 M DTT, 50 μl; 0.1 M PMSF, 100 μl; ddH_2_O, 7.25 ml) for 25 min at RT. Subsequently, the cells were centrifugated and resuspended in 100 mM sucrose and spread on slides with 1% PFA (pH 9.2) containing 0.15% Triton X-100. Slides were then placed in a humidified chamber for at least 3 hours. Last, slides were washed twice for 3 min in 0.4% Photoflo (Kodak) and air-dried at RT. Slides were either used for immunostaining immediately or stored at -80°C. The immunostaining procedure was conducted as described above, and details of primary and secondary antibodies are listed in supplementary table S2. Spermatocytes were staged according to previously published papers(*23*).

### Single-cell RNA sequencing (scRNA-seq) analysis

Samples from adult *Shoc1* KI and littermate control mice were used for scRNA-seq. Detailed scRNA-seq analysis of testicular samples has been described in the previous study(*49*). Briefly, individual mouse testicular cells were obtained through enzymatic digestion and loaded on a 10 × Genomics chip to form gel beads in emulsion, which were subjected to reverse transcription and PCR amplification, and then sequenced on a Illumina NovaSeq 6000. A cell-gene expression matrix was obtained with Cell Ranger software (10 × Genomics). After quality control, standardization and dimensionality reduction, cell identification and clustering analysis were performed in the R environment. The Seurat *FindMarkers* function was used for identification of DEGs in the *Shoc1* KI versus control pachytene cluster groups. Gene Ontology (GO) biological processes, Kyoto Encyclopedia of Genes and Genomes (KEGG) pathways, and GSEA were performed using WebGestalt 2024 (https://www.webgestalt.org).

### Purification of pachytene spermatocytes

The modified STA-PUT velocity sedimentation was used for purification of pachytene spermatocytes from mouse testicular tissues as previously reported(*49, 50*). To obtain testicular cell suspension, mouse testes were enzymatically digested with 4 mg/ml collagenase type IV (17104-019, Life Technologies), 2.5 mg/ml hyaluronidase (H3506, Sigma-Aldrich), and 1 mg/ml trypsin (T8003, Sigma-Aldrich) at 37°C for 20 min. Next, specific cell populations are enriched by gravity sedimentation through a discontinuous bovine serum albumin (BSA) density gradient. The cell fractions are then manually collected and the purity of different types of cells were measured by IF staining and morphology. Each purification was performed on testes from four mice. The spermatocyte samples with a purity of more than 75% were used for subsequent sequencing.

### Hi-C library construction

Briefly, isolated cells were fixed with 2% formaldehyde at RT for 10 min. Then formaldehyde was quenched with glycine for 10 min at RT. Cells were washed with 1 × PBS for two times and then lysed in 50 μl lysis buffer (10 mM Tris-HCl pH 7.4, 10 mM NaCl, 0.1 mM EDTA, 0.5% NP-40 and proteinase inhibitor cocktail) on ice for 50 min. After centrifugation at 3000 rpm for 5 min at 4L, the supernatant was discarded with a pipette carefully. Chromatin was solubilized in 0.5% SDS and incubated at 62L for 10 min. SDS was quenched by 10% Triton X-100 at 37L for 30 min. Then the nuclei were digested with 50 U DpnII at 37L overnight with rotation. DpnII was then inactivated at 62L for 20 min. To incorporate biotin-labeled dCTP into DNA, dATP, dGTP, dTTP, biotin-14-dCTP and Klenow were added to the solution and the reaction was carried out at 37L for 1.5 h with rotation. The fragments were ligated at RT for 6 h with rotation. This was followed by reversal of crosslink and DNA purification. DNA was sheared to 300-500 bp with Covaris M220. The biotin-labeled DNA was then pulled down with 10 μl Dynabeads MyOne Streptavidin C1 (Invitrogen, 65001). Sequencing library preparation was performed on beads, including end-repair, dATP tailing and adaptor-ligation. DNA was eluted twice by adding 20 μl water to the tube and incubation at 66L for 20 min. 9-15 cycles of PCR amplification were performed with Extaq (Takara, RR001). Finally, size selection was done with AMPure XP beads and fragments ranging from 200-1000 bp were selected. The library was sequenced on MGI DNBSEQ T7 according to the manufacturer’s instruction.

### Hi-C data analysis

We obtained WT Hi-C dataset from mouse cells at the pachytene stage from published work (accession no. GSE109344)(*28*). Fastq files of Hi-C libraries were processed by the Juicer pipeline with default settings (v1.6)(*51*), and the mm10 reference genome was used for mapping. The obtained .hic files were converted into cool format using the hic2cool package for subsequent downstream analyses. The cis and trans expected contact probability was calculated using the cooltools package (v0.7.0)(*52*). Contact probability (P(s)) curves were computed from the cool files binned at 5 kb resolution using cooltools.expected_cis from Cooltools. For compartment analysis, eigenvector decomposition was performed at a resolution of 500 Kb using the eigenvector function in Juicer. The TADs insulation score was calculated at 25 Kb resolution by Cooltools. The pileups of 500 kb genomic regions flanking the TADs boundaries were performed at 25 kb resolution using coolpup.py under the local mode (https://github.com/open2c/coolpuppy, v1.1.0)(*53*). The identification of COs-DSBs regions follows the method described in the previously published study(*29*). Specifically, the positions of DSB hotspots were obtained from the previously published DMC1 ChIP-Seq data(*54*) and converted into 10 kb genomic bins. The positions of CO hotspots were obtained from the single sperm DNA sequencing data(*55*). The 10 kb genomic bins containing DSBs were designated as COs-DSBs if they overlapped with CO hotspots. Pileup heatmaps for 2 Mb genomic regions centered at the selected COs-DSBs regions were generated at a resolution of 5 kb. The observed/expected (O/E) interaction values were extracted and statistically summarized in Python to compare the chromatin interactions around COs-DSBs.

### Statistical analysis

The data are shown as the mean ± SEM unless otherwise indicated. Statistical significance was determined using the Student’s t-test (for the results in Figures 3M,N,O, 4B,D,F,H,J,L,O,P,Q,R and 6F; Figures S5B, S6B, S7B and S8B,D,F,H and S10B,D; unpaired, two-tailed), or one-way ANOVA of variance (for the results in Figure 3D) with GraphPad Prism 9 software. The confidence interval was 95%. Statistical significance was determined using the Student’s t-test (for the results in Figures 5M, and 7G; unpaired, two-tailed) with R software (version 4.3.1; R Foundation for Statistical Computing, Vienna, Austria). *P* < 0.05 was considered statistically significant. Statistical parameters are reported in the figures or their legends.

## SUPPLEMENTARY MATERIALS

Fig. S1. Specific regions of human and mouse SHOC1 required for interaction with binding partners.

Fig. S2. Effects of SHOC1 variants assessed by in vitro analyses.

Fig. S3. Meiotic arrest phenotype in the patient carrying the missense variant (p.Q590R) within the XPF-like domain in *SHOC1*.

Fig. S4. *Shoc1* KI caused follicular dysplasia and female infertility.

Fig. S5. Meiotic prophase I analysis.

Fig. S6. Visualization of both human and mouse SHOC1 mutants through molecular docking.

Fig. S7. DSBs repair defects in Shoc1 KI spermatocytes.

Fig. S8. Isolation of pachytene spermatocytes by modified STA-PUT.

Fig. S9. *Shoc1* KI induced excessive accumulation of DNA damage response (DDR) factors and establishment of MSUC on autosomes.

Table S1. Information of the primers.

Table S2. Information of the antibodies.

Table S3. Clinical and semen characteristics in Chinese men with bi-allelic *SHOC1* variants

## Acknowledgments

We thank Professor Qinghua Shi for providing REDIC1 and M1AP antibodies. We thank Professor Chao Yu for providing TEX11 antibody. We thank Professor Hongbin Liu for providing HEI10 antibody. We thank Professor Kui Liu for providing SYCP1 antibody. We are grateful to all patients and their family members for participating in our research.

## Author contributors

Z.Y.X., S.Q., and J.Z.Y. contributed equally to this work. L.P., Z.Z., L.Z., and Y.C.C. conceived and designed the study. Z.Y.X. performed most experiments and analyzed the data. Data analysis and manuscript writing were conducted by Z.Y.X., S.Q., and Y.C.C., L.P., Z.Z., L.Z., and Y.C.C. reviewed and revised the manuscript. J.Z.Y., Z.J.P., and X.S. assisted with H&E and IF assays, L.N., S.Y.F., and B.H.W., and G.Y. with WB and co-IP, Q.D.W., H.S., and B.X.J. with plasmid construction, Z.E.L., and H.Y.H., and Y.C. with spermatocyte spread, Z.Y.C., C.H.X., Z.J., T.R.H., L.J.X., and Z.F.J. with collection of human testicular tissues and blood samples. Z.Y.X., and X.S., and Z.L.Y. analyzed single-cell RNA-seq data. Z.Y.X., and S.Q. performed bioinformatic analyses. All authors reviewed and approved the final version. L.P., Z.Z., L.Z., Y.C.C., and Z.Y.X. verified the raw data.

## Data and materials availability

The scRNA-seq matrix data generated in this study has been deposited in Dryad with accession number: 10.5061/dryad,15dv41p7t. The Hi-C data generated in this study has been deposited in Dryad with accession number: 10.5061/dryad.3n5tb2rvx. Control Hi-C dataset from mouse cells at the pachytene stage was obtained from published work (accession no. GSE109344)(*28*). All other relevant data supporting the key findings of this study are available within the article and its supplementary information files or from the corresponding authors upon reasonable request. Source data are provided with this paper. The data supporting this study are available from the corresponding author upon reasonable request.

## REFERENCES AND NOTES

1. A. Agarwal, S. Baskaran, N. Parekh, C. L. Cho, R. Henkel, S. Vij, M. Arafa, M. K. Panner Selvam, R. Shah, Male infertility. Lancet 397, 319–333 (2021).

2. S. Minhas, C. Bettocchi, L. Boeri, P. Capogrosso, J. Carvalho, N. C. Cilesiz, A. Cocci, G. Corona, K. Dimitropoulos, M. Gul, G. Hatzichristodoulou, T. H. Jones, A. Kadioglu, J. I. Martinez Salamanca, U. Milenkovic, V. Modgil, G. I. Russo, E. C. Serefoglu, T. Tharakan, P. Verze, A. Salonia, E. A. U. W. G. o. M. Sexual, H. Reproductive, European Association of Urology Guidelines on Male Sexual and Reproductive Health: 2021 Update on Male Infertility. Eur Urol 80, 603–620 (2021).

3. H. Tournaye, C. Krausz, R. D. Oates, Novel concepts in the aetiology of male reproductive impairment. Lancet Diabetes Endocrinol 5, 544–553 (2017).

4. S. Y. Jiao, Y. H. Yang, S. R. Chen, Molecular genetics of infertility: loss-of-function mutations in humans and corresponding knockout/mutated mice. Hum Reprod Update 27, 154–189 (2021).

5. Y. Fujiwara, Y. Horisawa-Takada, E. Inoue, N. Tani, H. Shibuya, S. Fujimura, R. Kariyazono, T. Sakata, K. Ohta, K. Araki, Y. Okada, K. I. Ishiguro, Meiotic cohesins mediate initial loading of HORMAD1 to the chromosomes and coordinate SC formation during meiotic prophase. PLoS Genet 16, e1009048 (2020).

6. T. Bhattacharyya, M. Walker, N. R. Powers, C. Brunton, A. D. Fine, P. M. Petkov, M. A. Handel, Prdm9 and Meiotic Cohesin Proteins Cooperatively Promote DNA Double-Strand Break Formation in Mammalian Spermatocytes. Curr Biol 29, 1002–1018 e1007 (2019).

7. J. H. Joo, S. Hong, M. T. Higashide, E. H. Choi, S. Yoon, M. S. Lee, H. A. Kang, A. Shinohara, N. Kleckner, K. P. Kim, RPA interacts with Rad52 to promote meiotic crossover and noncrossover recombination. Nucleic Acids Res 52, 3794–3809 (2024).

8. A. Pyatnitskaya, V. Borde, A. De Muyt, Crossing and zipping: molecular duties of the ZMM proteins in meiosis. Chromosoma 128, 181–198 (2019).

9. L. Payero, E. Alani, Crossover recombination between homologous chromosomes in meiosis: recent progress and remaining mysteries. Trends Genet 41, 47–59 (2025).

10. T. Wang, H. Wang, Q. Lian, Q. Jia, C. You, G. P. Copenhaver, C. Wang, Y. Wang, HEI10 is subject to phase separation and mediates RPA1a degradation during meiotic interference-sensitive crossover formation. Proc Natl Acad Sci U S A 120, e2310542120 (2023).

11. H. Qiao, H. B. Prasada Rao, Y. Yang, J. H. Fong, J. M. Cloutier, D. C. Deacon, K. E. Nagel, R. K. Swartz, E. Strong, J. K. Holloway, P. E. Cohen, J. Schimenti, J. Ward, N. Hunter, Antagonistic roles of ubiquitin ligase HEI10 and SUMO ligase RNF212 regulate meiotic recombination. Nat Genet 46, 194–199 (2014).

12. P. R. Chua, G. S. Roeder, Zip2, a meiosis-specific protein required for the initiation of chromosome synapsis. Cell 93, 349–359 (1998).

13. Q. Zhang, S. Y. Ji, K. Busayavalasa, C. Yu, SPO16 binds SHOC1 to promote homologous recombination and crossing-over in meiotic prophase I. Sci Adv 5, eaau9780 (2019).

14. Q. Zhang, J. Shao, H. Y. Fan, C. Yu, Evolutionarily-conserved MZIP2 is essential for crossover formation in mammalian meiosis. Commun Biol 1, 147 (2018).

15. N. Macaisne, M. Novatchkova, L. Peirera, D. Vezon, S. Jolivet, N. Froger, L. Chelysheva, M. Grelon, R. Mercier, SHOC1, an XPF endonuclease-related protein, is essential for the formation of class I meiotic crossovers. Curr Biol 18, 1432–1437 (2008).

16. W. Wang, L. Meng, J. He, L. Su, Y. Li, C. Tan, X. Xu, H. Nie, H. Zhang, J. Du, G. Lu, M. Luo, G. Lin, C. Tu, Y. Q. Tan, Bi-allelic variants in SHOC1 cause non-obstructive azoospermia with meiosis arrest in humans and mice. Mol Hum Reprod 28, (2022).

17. M. F. Guiraldelli, A. Felberg, L. P. Almeida, A. Parikh, R. O. de Castro, R. J. Pezza, SHOC1 is a ERCC4-(HhH)2-like protein, integral to the formation of crossover recombination intermediates during mammalian meiosis. PLoS Genet 14, e1007381 (2018).

18. M. Shinohara, S. D. Oh, N. Hunter, A. Shinohara, Crossover assurance and crossover interference are distinctly regulated by the ZMM proteins during yeast meiosis. Nat Genet 40, 299–309 (2008).

19. E. Cannavo, A. Sanchez, R. Anand, L. Ranjha, J. Hugener, C. Adam, A. Acharya, N. Weyland, X. Aran-Guiu, J. B. Charbonnier, E. R. Hoffmann, V. Borde, J. Matos, P. Cejka, Regulation of the MLH1-MLH3 endonuclease in meiosis. Nature 586, 618–622 (2020).

20. A. N. Yatsenko, A. P. Georgiadis, A. Ropke, A. J. Berman, T. Jaffe, M. Olszewska, B. Westernstroer, J. Sanfilippo, M. Kurpisz, A. Rajkovic, S. A. Yatsenko, S. Kliesch, S. Schlatt, F. Tuttelmann, X-linked TEX11 mutations, meiotic arrest, and azoospermia in infertile men. N Engl J Med 372, 2097–2107 (2015).

21. F. Yang, S. Silber, N. A. Leu, R. D. Oates, J. D. Marszalek, H. Skaletsky, L. G. Brown, S. Rozen, D. C. Page, P. J. Wang, TEX11 is mutated in infertile men with azoospermia and regulates genome-wide recombination rates in mouse. EMBO Mol Med 7, 1198–1210 (2015).

22. Y. Li, Y. Wu, I. Khan, J. Zhou, Y. Lu, J. Ye, J. Liu, X. Xie, C. Hu, H. Jiang, S. Fan, H. Zhang, Y. Zhang, X. Jiang, B. Xu, H. Ma, Q. Shi, M1AP interacts with the mammalian ZZS complex and promotes male meiotic recombination. EMBO Rep 24, e55778 (2023).

23. S. Fan, Y. Wang, H. Jiang, X. Jiang, J. Zhou, Y. Jiao, J. Ye, Z. Xu, Y. Wang, X. Xie, H. Zhang, Y. Li, W. Liu, X. Zhang, H. Ma, B. Shi, Y. Zhang, M. Zubair, W. Shah, Z. Xu, B. Xu, Q. Shi, A novel recombination protein C12ORF40/REDIC1 is required for meiotic crossover formation. Cell Discov 9, 88 (2023).

24. J. K. Holloway, J. Booth, W. Edelmann, C. H. McGowan, P. E. Cohen, MUS81 generates a subset of MLH1-MLH3-independent crossovers in mammalian meiosis. PLoS Genet 4, e1000186 (2008).

25. S. D. Desjardins, J. Simmonds, I. Guterman, K. Kanyuka, A. J. Burridge, A. J. Tock, E. Sanchez-Moran, F. C. H. Franklin, I. R. Henderson, K. J. Edwards, C. Uauy, J. D. Higgins, FANCM promotes class I interfering crossovers and suppresses class II non-interfering crossovers in wheat meiosis. Nat Commun 13, 3644 (2022).

26. C. Yao, C. Yang, L. Zhao, P. Li, R. Tian, H. Chen, Y. Guo, Y. Huang, E. Zhi, J. Zhai, H. Sun, J. Zhang, Y. Hong, L. Zhang, Z. Ji, F. Zhang, Z. Zhou, Z. Li, Bi-allelic SHOC1 loss-of-function mutations cause meiotic arrest and non-obstructive azoospermia. J Med Genet 58, 679–686 (2021).

27. H. Ma, T. Li, X. Xie, L. Jiang, J. Ye, C. Gong, H. Jiang, S. Fan, H. Zhang, B. Shi, B. Zhang, X. Jiang, Y. Li, J. Zhou, J. Xu, X. Zhang, X. Hou, H. Yin, Y. Zhang, Q. Shi, RAD51AP2 is required for efficient meiotic recombination between X and Y chromosomes. Sci Adv 8, eabk1789 (2022).

28. Y. Wang, H. Wang, Y. Zhang, Z. Du, W. Si, S. Fan, D. Qin, M. Wang, Y. Duan, L. Li, Y. Jiao, Y. Li, Q. Wang, Q. Shi, X. Wu, W. Xie, Reprogramming of Meiotic Chromatin Architecture during Spermatogenesis. Mol Cell 73, 547–561 e546 (2019).

29. W. Zuo, G. Chen, Z. Gao, S. Li, Y. Chen, C. Huang, J. Chen, Z. Chen, M. Lei, Q. Bian, Stage-resolved Hi-C analyses reveal meiotic chromosome organizational features influencing homolog alignment. Nat Commun 12, 5827 (2021).

30. G. Cheng, F. Pratto, K. Brick, X. Li, B. Alleva, M. Huang, G. Lam, R. D. Camerini-Otero, High resolution maps of chromatin reorganization through mouse meiosis reveal novel features of the 3D meiotic structure. bioRxiv, (2024).

31. Z. Luo, X. Wang, H. Jiang, R. Wang, J. Chen, Y. Chen, Q. Xu, J. Cao, X. Gong, J. Wu, Y. Yang, W. Li, C. Han, C. Y. Cheng, M. G. Rosenfeld, F. Sun, X. Song, Reorganized 3D Genome Structures Support Transcriptional Regulation in Mouse Spermatogenesis. iScience 23, 101034 (2020).

32. L. Patel, R. Kang, S. C. Rosenberg, Y. Qiu, R. Raviram, S. Chee, R. Hu, B. Ren, F. Cole, K. D. Corbett, Dynamic reorganization of the genome shapes the recombination landscape in meiotic prophase. Nat Struct Mol Biol 26, 164–174 (2019).

33. K. G. Alavattam, S. Maezawa, P. R. Andreassen, S. H. Namekawa, Meiotic sex chromosome inactivation and the XY body: a phase separation hypothesis. Cell Mol Life Sci 79, 18 (2021).

34. J. M. Turner, Meiotic Silencing in Mammals. Annu Rev Genet 49, 395–412 (2015).

35. J. Zhang, M. Gurusaran, Y. Fujiwara, K. Zhang, M. Echbarthi, E. Vorontsov, R. Guo, D. F. Pendlebury, I. Alam, G. Livera, M. Emmanuelle, P. J. Wang, J. Nandakumar, O. R. Davies, H. Shibuya, The BRCA2-MEILB2-BRME1 complex governs meiotic recombination and impairs the mitotic BRCA2-RAD51 function in cancer cells. Nat Commun 11, 2055 (2020).

36. L. Bai, P. Li, Y. Xiang, X. Jiao, J. Chen, L. Song, Z. Liang, Y. Liu, Y. Zhu, L. Y. Lu, BRCA1 safeguards genome integrity by activating chromosome asynapsis checkpoint to eliminate recombination-defective oocytes. Proc Natl Acad Sci U S A 121, e2401386121 (2024).

37. K. Arora, K. D. Corbett, The conserved XPF:ERCC1-like Zip2:Spo16 complex controls meiotic crossover formation through structure-specific DNA binding. Nucleic Acids Res 47, 2365-2376 (2019).

38. A. De Muyt, A. Pyatnitskaya, J. Andreani, L. Ranjha, C. Ramus, R. Laureau, A. Fernandez-Vega, D. Holoch, E. Girard, J. Govin, R. Margueron, Y. Coute, P. Cejka, R. Guerois, V. Borde, A meiotic XPF-ERCC1-like complex recognizes joint molecule recombination intermediates to promote crossover formation. Genes Dev 32, 283–296 (2018).

39. C. Krausz, A. Riera-Escamilla, D. Moreno-Mendoza, K. Holleman, F. Cioppi, F. Algaba, M. Pybus, C. Friedrich, M. J. Wyrwoll, E. Casamonti, S. Pietroforte, L. Nagirnaja, A. M. Lopes, S. Kliesch, A. Pilatz, D. T. Carrell, D. F. Conrad, E. Ars, E. Ruiz-Castane, K. I. Aston, W. M. Baarends, F. Tuttelmann, Genetic dissection of spermatogenic arrest through exome analysis: clinical implications for the management of azoospermic men. Genet Med 22, 1956–1966 (2020).

40. F. Yang, K. Gell, G. W. van der Heijden, S. Eckardt, N. A. Leu, D. C. Page, R. Benavente, C. Her, C. Hoog, K. J. McLaughlin, P. J. Wang, Meiotic failure in male mice lacking an X-linked factor. Genes Dev 22, 682–691 (2008).

41. P. Benn, Trisomy 16 and trisomy 16 Mosaicism: a review. Am J Med Genet 79, 121–133 (1998).

42. T. G. Cooper, E. Noonan, S. von Eckardstein, J. Auger, H. W. Baker, H. M. Behre, T. B. Haugen, T. Kruger, C. Wang, M. T. Mbizvo, K. M. Vogelsong, World Health Organization reference values for human semen characteristics. Hum Reprod Update 16, 231–245 (2010).

43. Y. Zhang, N. Li, Z. Ji, H. Bai, N. Ou, R. Tian, P. Li, E. Zhi, Y. Huang, J. Zhao, Y. Han, J. Zhang, Y. Zhou, Z. Li, C. Yao, Bi-allelic MEI1 variants cause meiosis arrest and non-obstructive azoospermia. J Hum Genet 68, 383–392 (2023).

44. S. Xu, J. Zhao, F. Gao, Y. Zhang, J. Luo, C. Zhang, R. Tian, E. Zhi, J. Zhang, F. Bai, H. Sun, F. Zhao, Y. Huang, P. Li, L. Jiang, Z. Li, C. Yao, Z. Zhou, A bi-allelic REC114 loss-of-function variant causes meiotic arrest and nonobstructive azoospermia. Clin Genet 105, 440–445 (2024).

45. H. Li, R. Durbin, Fast and accurate short read alignment with Burrows-Wheeler transform. Bioinformatics 25, 1754–1760 (2009).

46. Y. Yan, H. Tao, J. He, S. Y. Huang, The HDOCK server for integrated protein-protein docking. Nat Protoc 15, 1829–1852 (2020).

47. H. Yang, C. Zhou, A. Dhar, N. P. Pavletich, Mechanism of strand exchange from RecA-DNA synaptic and D-loop structures. Nature 586, 801–806 (2020).

48. N. Ou, Y. Wang, S. Xu, J. Luo, C. Zhang, Y. Zhang, X. Shi, M. Xiong, L. Zhao, Z. Ji, Y. Zhang, J. Zhao, H. Bai, R. Tian, P. Li, E. Zhi, Y. Huang, W. Chen, R. Wang, Y. Jin, D. Wang, Z. Li, H. Chen, C. Yao, Primate-Specific DAZ Regulates Translation of Cell Proliferation-Related mRNAs and is Essential for Maintenance of Spermatogonia. Adv Sci (Weinh) 11, e2400692 (2024).

49. L. Zhao, C. Yao, X. Xing, T. Jing, P. Li, Z. Zhu, C. Yang, J. Zhai, R. Tian, H. Chen, J. Luo, N. Liu, Z. Deng, X. Lin, N. Li, J. Fang, J. Sun, C. Wang, Z. Zhou, Z. Li, Single-cell analysis of developing and azoospermia human testicles reveals central role of Sertoli cells. Nat Commun 11, 5683 (2020).

50. M. Da Ros, T. Lehtiniemi, O. Olotu, O. Meikar, N. Kotaja, Enrichment of Pachytene Spermatocytes and Spermatids from Mouse Testes Using Standard Laboratory Equipment. J Vis Exp, (2019).

51. N. C. Durand, M. S. Shamim, I. Machol, S. S. Rao, M. H. Huntley, E. S. Lander, E. L. Aiden, Juicer Provides a One-Click System for Analyzing Loop-Resolution Hi-C Experiments. Cell Syst 3, 95–98 (2016).

52. Open2C, N. Abdennur, S. Abraham, G. Fudenberg, I. M. Flyamer, A. A. Galitsyna, A. Goloborodko, M. Imakaev, B. A. Oksuz, S. V. Venev, Y. Xiao, Cooltools: Enabling high-resolution Hi-C analysis in Python. PLoS Comput Biol 20, e1012067 (2024).

53. I. M. Flyamer, R. S. Illingworth, W. A. Bickmore, Coolpup.py: versatile pile-up analysis of Hi-C data. Bioinformatics 36, 2980–2985 (2020).

54. B. Davies, E. Hatton, N. Altemose, J. G. Hussin, F. Pratto, G. Zhang, A. G. Hinch, D. Moralli, D. Biggs, R. Diaz, C. Preece, R. Li, E. Bitoun, K. Brick, C. M. Green, R. D. Camerini-Otero, S. R. Myers, P. Donnelly, Re-engineering the zinc fingers of PRDM9 reverses hybrid sterility in mice. Nature 530, 171–176 (2016).

55. A. G. Hinch, G. Zhang, P. W. Becker, D. Moralli, R. Hinch, B. Davies, R. Bowden, P. Donnelly, Factors influencing meiotic recombination revealed by whole-genome sequencing of single sperm. Science 363, (2019).

