## Supplementary figures and images for "The XPF-like domain in SHOC1 required for homologous recombination and safeguarding autosome from meiotic silencing of unsynapsed chromatin"

### Figure S1

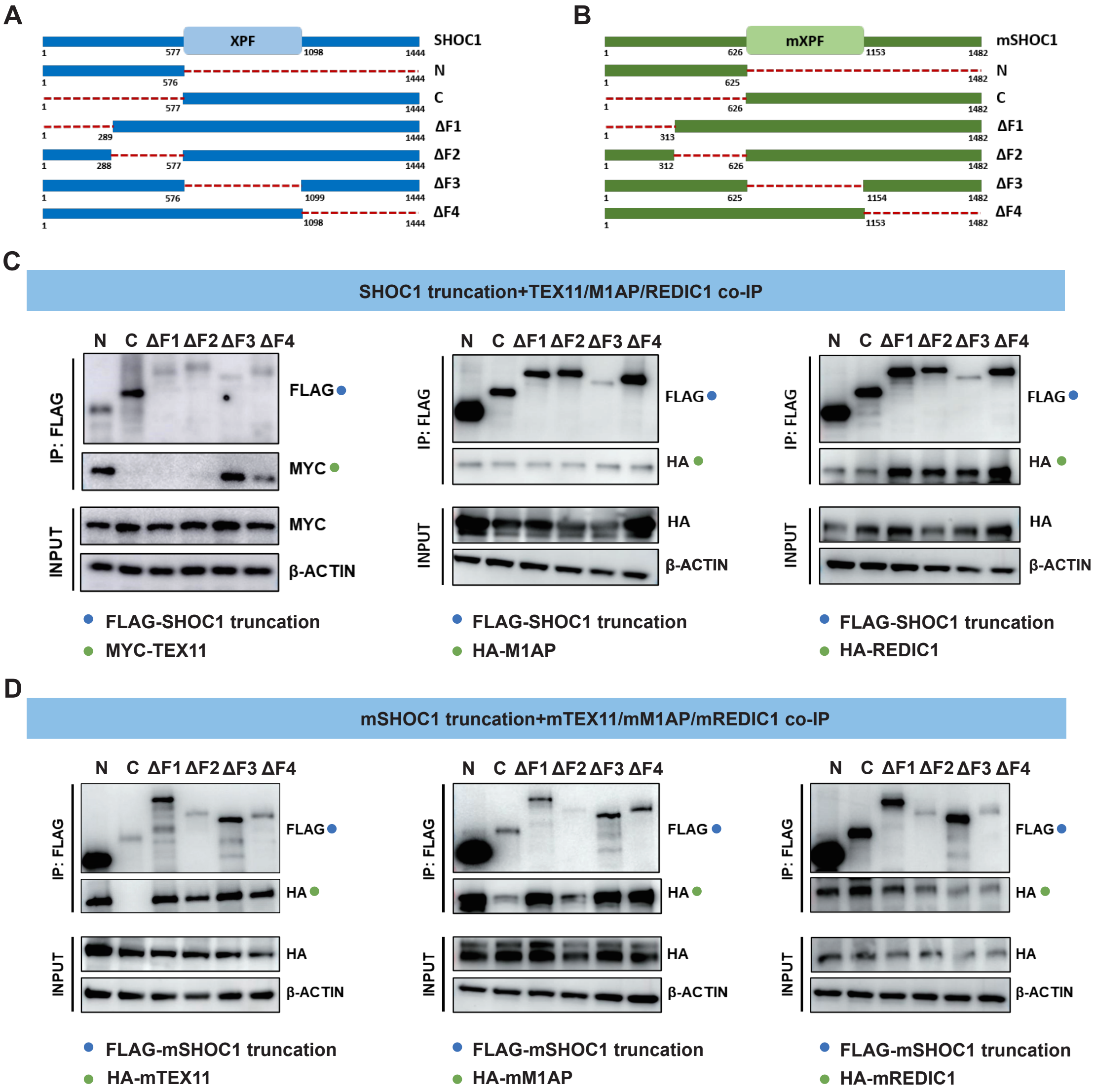

### Figure S2

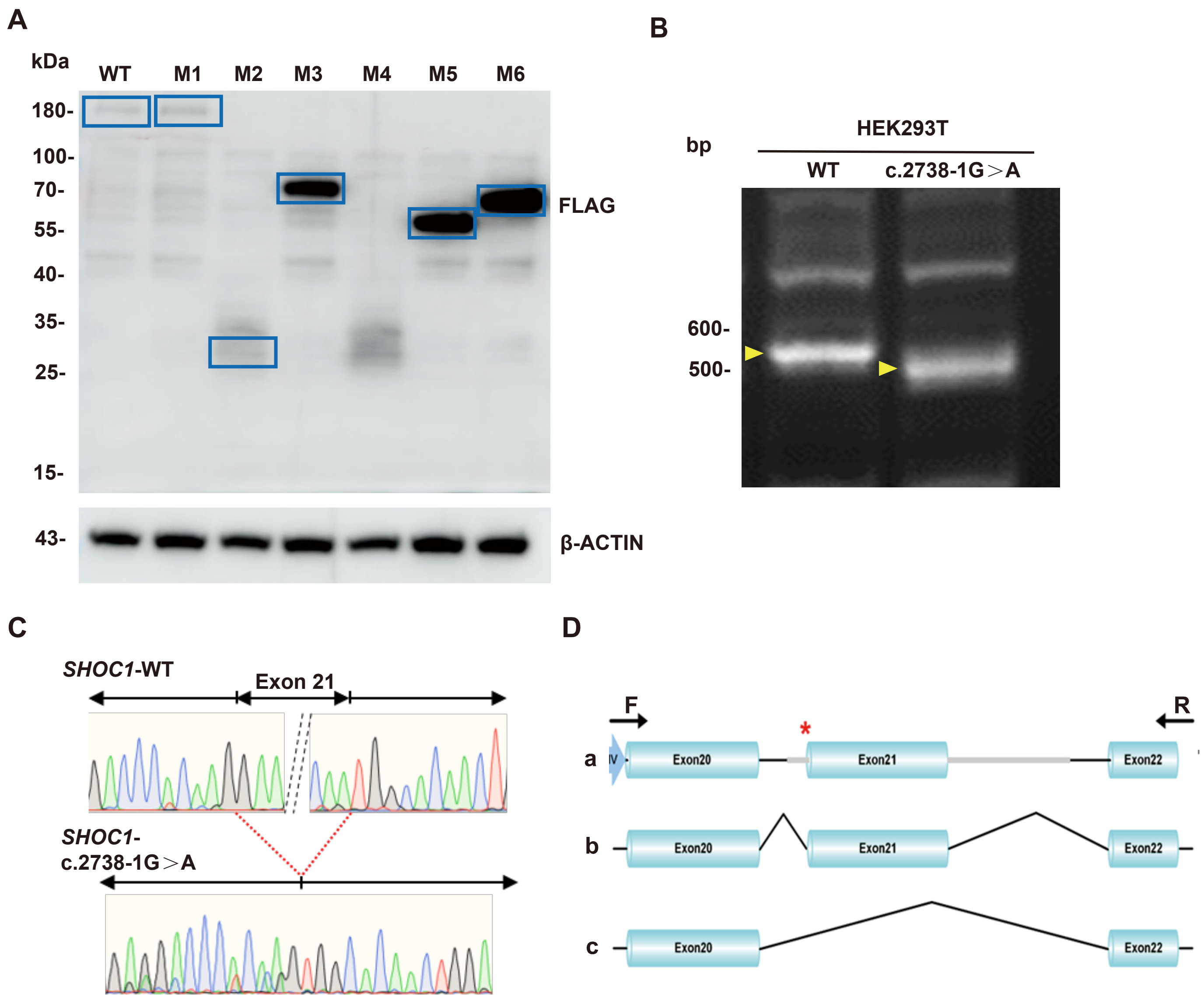

### Figure S3

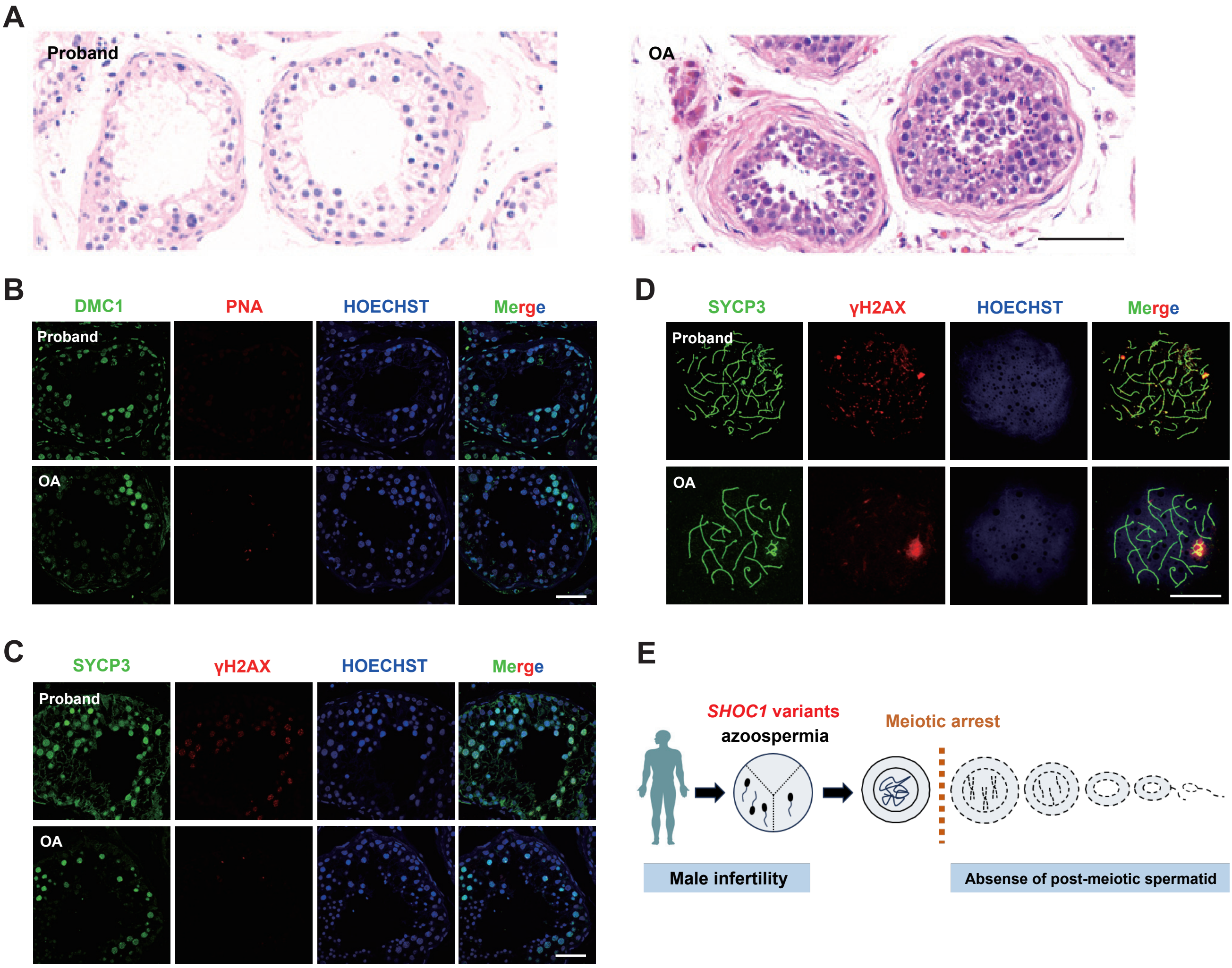

### Figure S4

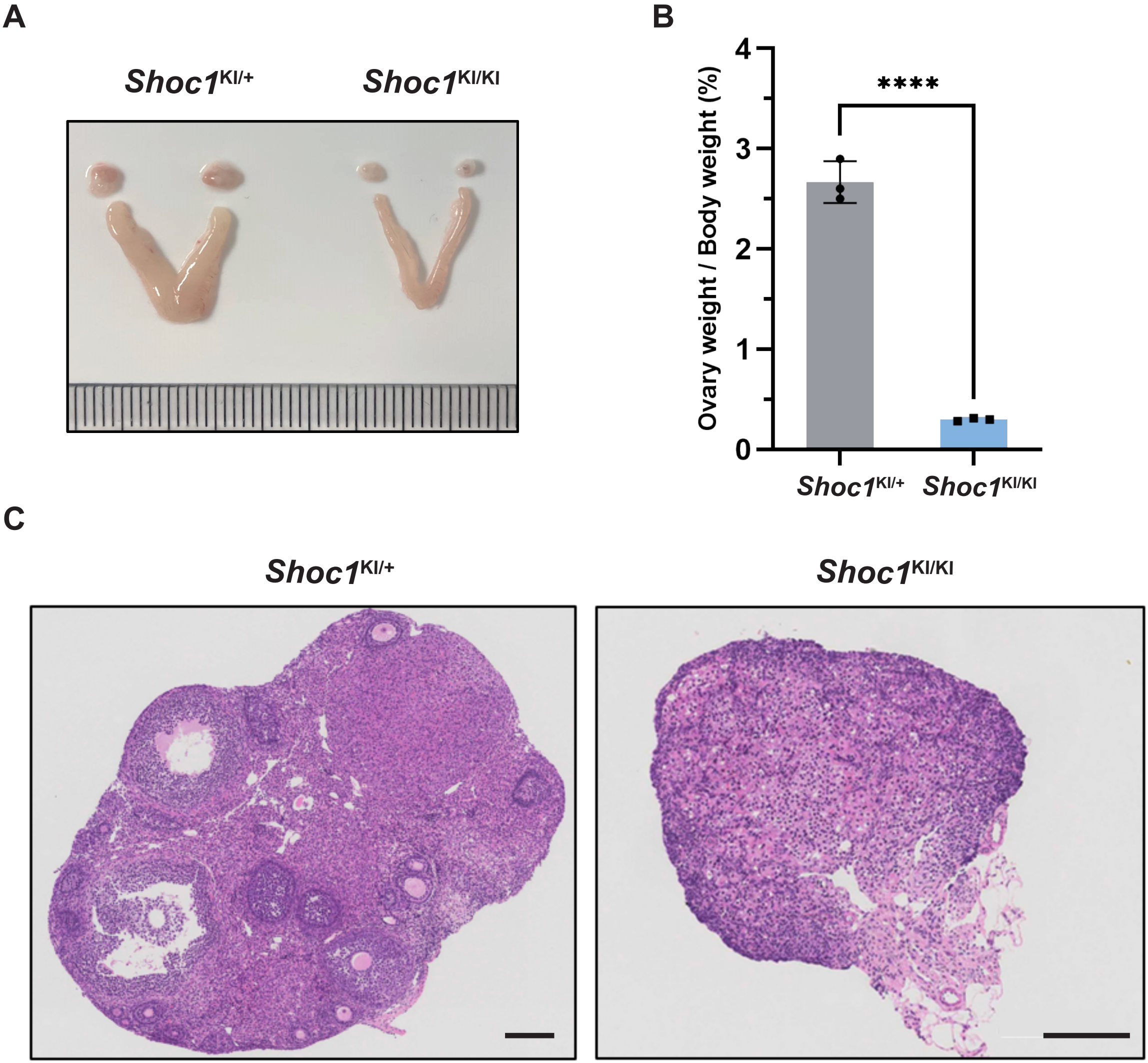

### Figure S5

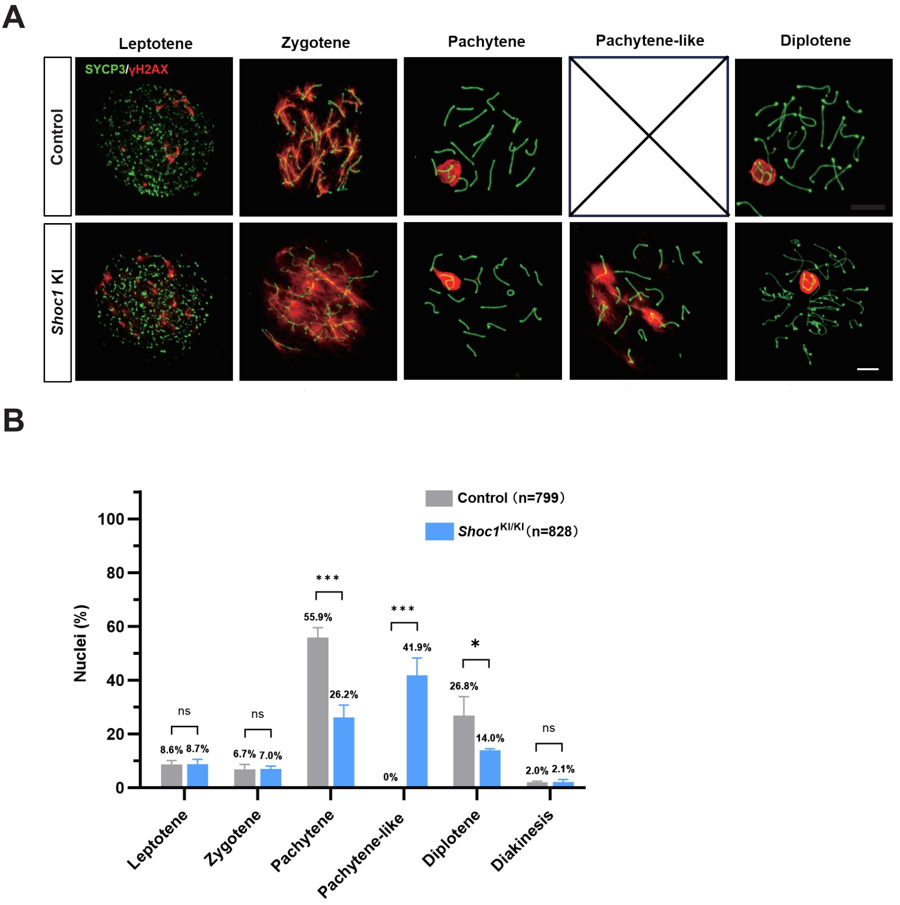

### Figure S6

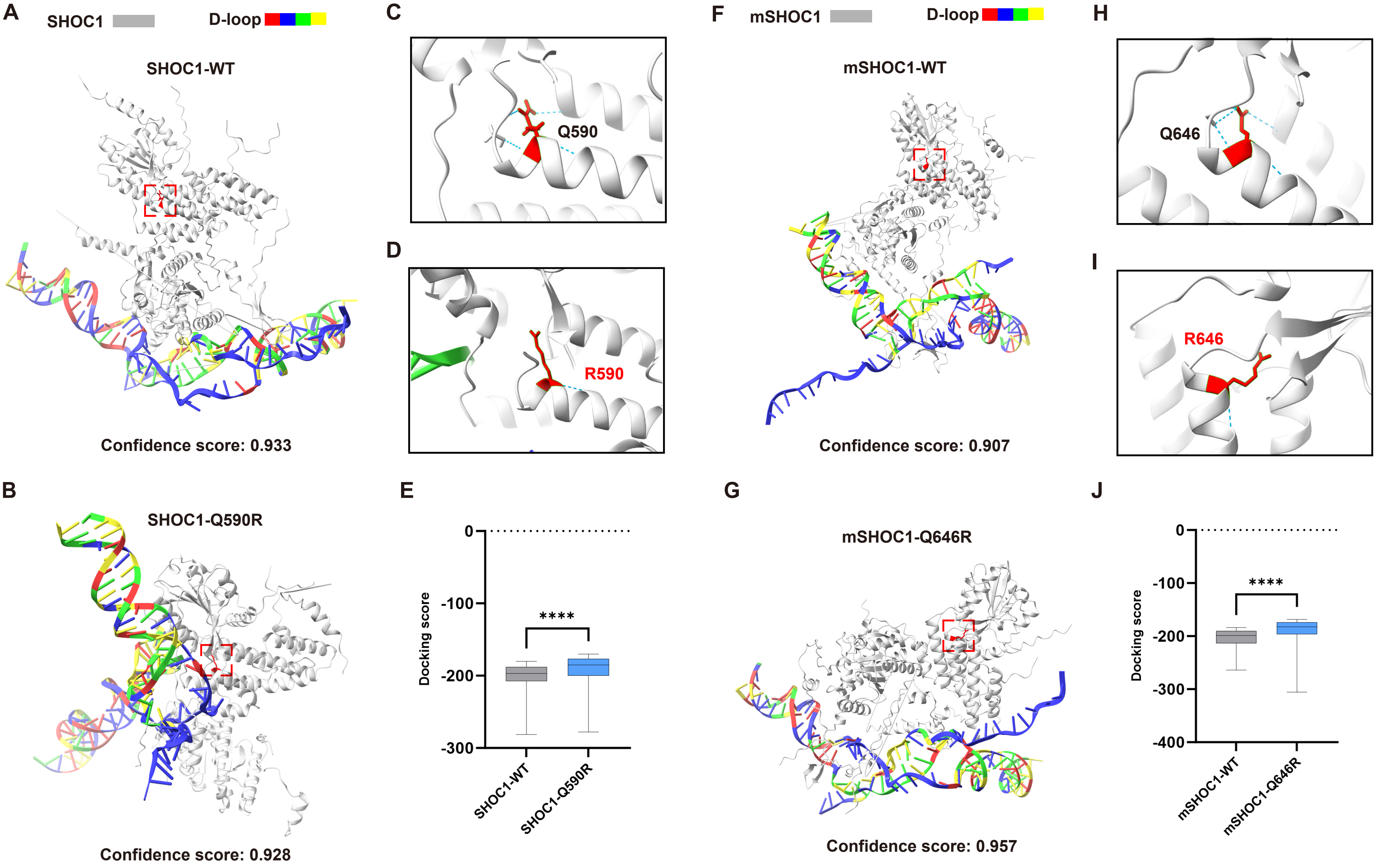

### Figure S7

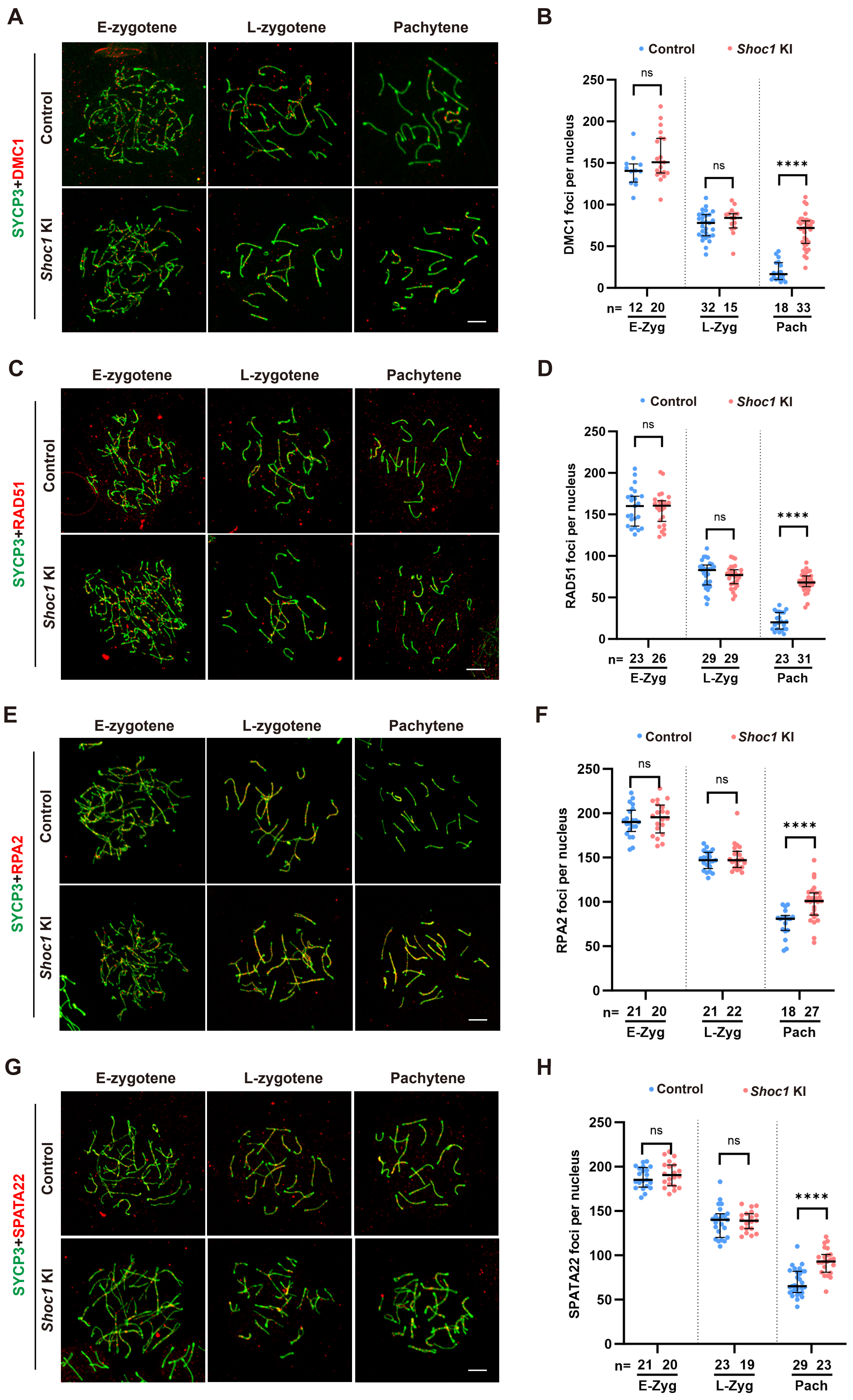

### Figure S8

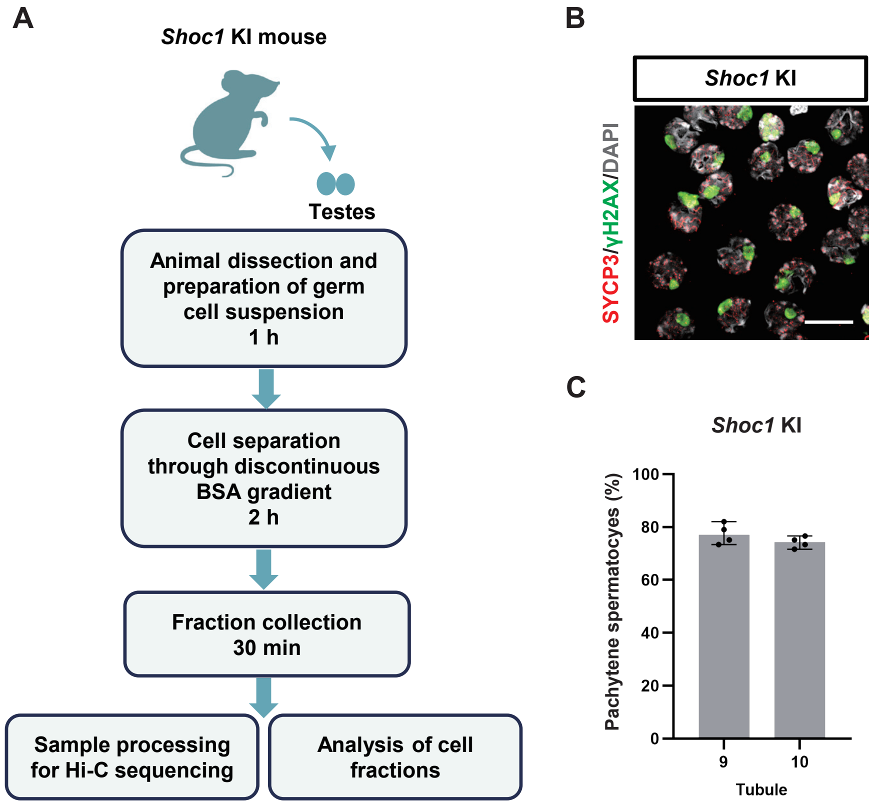

### Figure S9

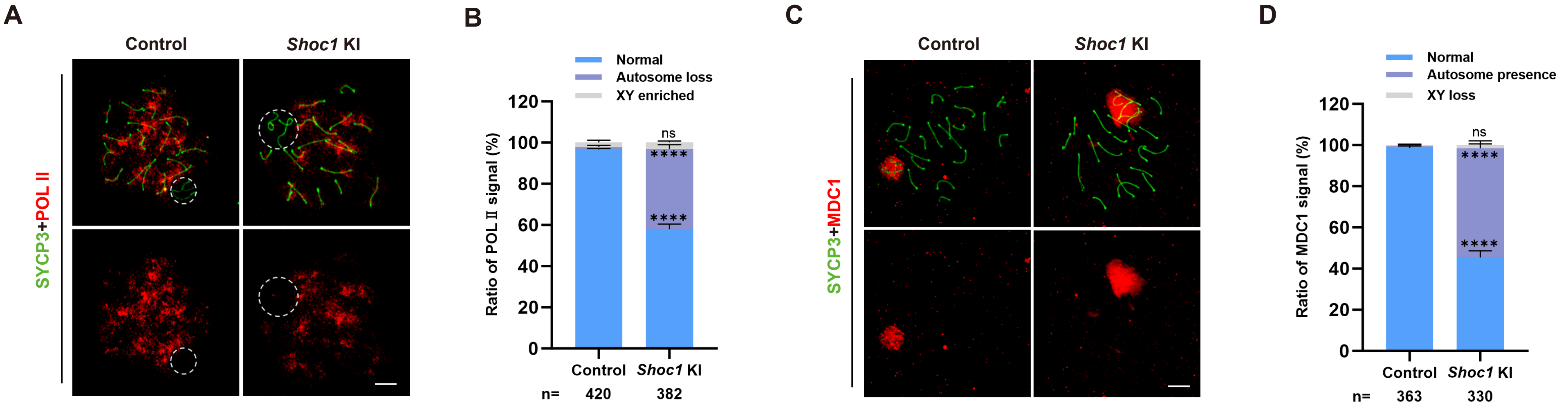
