## Supplementary material for "The XPF-like domain in SHOC1 required for homologous recombination and safeguarding autosome from meiotic silencing of unsynapsed chromatin": Figure legend

**Figure S1. Specific regions of human and mouse SHOC1 required for interaction with binding partners.**

**(A and B)** Schematic diagrams of truncated human (A) and mouse (B) SHOC1 protein constructs.

**(C)** Co-IP assays showing interactions between full-length human TEX11, M1AP, or REDIC1 (detected using N-terminal HA- or MYC-tags) and truncated human SHOC1 constructs (detected using an N-terminal FLAG-tag) in HEK293T cells. Input lysates were included.

**(D)** Co-IP assays demonstrating interactions between full-length mouse TEX11, M1AP, or REDIC1 (detected using N-terminal HA-tags) and truncated mouse SHOC1 constructs (detected using an N-terminal FLAG-tag) in HEK293T cells. Input lysates were included.

**Figure S2. Effects of *SHOC1* variants assessed by *in vitro* analyses.**

**(A)** WB analysis of 3×FLAG-SHOC1 fusion protein expression in HEK293T cells transfected with WT or mutant (M1 to M6) plasmids. Target protein bands are indicated by blue rectangles.

**(B)** Reverse transcription-PCR products of pcMINI vectors expressing WT or c. G2738-1A *SHOC1* minigenes (containing genomic sequences: intron [179 bp]-exon 21 [118 bp]-intron [112 bp]) in HEK293T cells.

**(C)** Sanger sequencing analysis of splicing products. WT transcripts exhibited canonical splicing, while mutant transcripts underwent aberrant splicing, resulting in exon 21 skipping.

**(D)** Schematic representation of canonical (WT) and aberrant (mutant) splicing, and predicted protein truncation caused by the c.G2738-1A variant in *SHOC1*. The red asterisk indicates the variant site.

**Figure S3. Meiotic arrest phenotype in the patient carrying the missense variant (p.Q590R) within the XPF-like domain in *SHOC1.***

**(A)** H&E staining of testicular sections from the proband (left) and an OA control (right). Scale bars, 50 μm.

**(B)** IF staining of testicular sections from the proband and an OA control with DMC1 (green) and PNA (red). Scale bars, 50 μm.

**(C)** IF staining of testicular sections from the proband and an OA control with SYCP3 (green) and γH2AX (red). Scale bars, 50 μm.

**(D)** Representative images of spread spermatocytes from the proband and an OA control co-stained with SYCP3 (green) and γH2AX (red). Scale bars, 20 μm.

**(E)** Schematic diagram of meiotic arrest and male infertility caused by bi-allelic *SHOC1* variants.

**Figure S4. *Shoc1* KI caused follicular dysplasia and female infertility.**

**(A)** Representative image comparing ovary size in adult *Shoc1* KI homozygous (*Shoc1*^KI/KI^) mice and littermate controls (*Shoc1*^KI/+^).

**(B)** Ovary-to-body weight ratios quantified for *Shoc1* KI homozygous (*Shoc1*^KI/KI^) mice and littermate controls (*Shoc1*^KI/+^) using two-tailed Student’s t-test; **** *P* < 0.0001; error bars, mean ± SEM.

**(C)** H&E staining of ovary sections from adult *Shoc1* KI homozygous (*Shoc1*^KI/KI^) mice and littermate controls (*Shoc1*^KI/+^). Scale bars, 100 μm.

**Figure S5.** **Meiotic prophase I analysis.**

**(A)** Representative images of spread spermatocytes from adult *Shoc1* KI homozygous (*Shoc1*^KI/KI^) mice and littermate controls (*Shoc1*^KI/+^) co-stained with SYCP3 (green) and γH2AX (red). Scale bars, 10 μm.

**(B)** Proportions of spermatocytes at defined substages using two-tailed Student’s t-test; *** *P* < 0.001; * *P* < 0.05; ns, not significant; error bars, mean ± SEM; n, the total number of nuclei analyzed.

**Figure S6. Visualization of both human and mouse SHOC1 mutants through molecular docking.**

**(A and B)** Structural model of WT (A) and mutant (B) human SHOC1 (predicted by AlphaFold3)/D-loop (PDB ID: 7JY7) complexes generated using the HDOCK server. Residue Q590 and its mutation R590 are highlighted in red dash rectangles, with hydrogen bond interactions indicated by blue dash lines. Confidence scores above 0.7 indicate high probability of binding.

**(C and D)** Zoom the view of Q590 (C) and its mutation R590 (D) residues.

**(E)** Comparison of docking scores between WT and mutant human SHOC1/D-loop complexes using two-tailed Student’s t-test, with lower (more negative) docking scores corresponding to increased binding energy; **** *P* < 0.0001; ns, error bars, mean ± SEM.

**(F and G)** Structural model of WT (F) and mutant (G) mouse SHOC1 (predicted by AlphaFold3)/D-loop (PDB ID: 7JY7) complexes generated using the HDOCK server. Residue Q646 and its mutation R646 are highlighted in red dash rectangles, with hydrogen bond interactions indicated by blue dash lines. Confidence scores above 0.7 indicate high probability of binding.

**(H and I)** Zoom the view of Q646 (H) and its mutation R646 (I) residues.

**(J)** Comparison of docking scores between WT and mutant mouse SHOC1/D-loop complexes using two-tailed Student’s t-test, with lower (more negative) docking scores corresponding to increased binding energy; **** *P* < 0.0001; ns, error bars, mean ± SEM.

**Figure S7. DSBs repair defects in *Shoc1* KI spermatocytes.**

**(A)** Representative images of spread spermatocytes from adult *Shoc1* KI homozygous (*Shoc1*^KI/KI^) mice and littermate controls (*Shoc1*^KI/+^) co-stained with SYCP3 (green) and DMC1 (red). Scale bars, 10 μm.

**(B)** Quantification of DMC1 foci per cell at indicated meiotic stages using two-tailed Student’s t-test; **** *P* < 0.0001; ns, not significant; error bars, mean ± SEM; n, the total number of nuclei analyzed.

**(C)** Representative images of spread spermatocytes from adult *Shoc1* KI homozygous (*Shoc1*^KI/KI^) mice and littermate controls (*Shoc1*^KI/+^) co-stained with SYCP3 (green) and RAD51 (red). Scale bars, 10 μm.

**(D)** Quantification of RAD51 foci per cell at indicated meiotic stages using two-tailed Student’s t-test; **** *P* < 0.0001; ns, not significant; error bars, mean ± SEM; n, the total number of nuclei analyzed.

**(E)** Representative images of spread spermatocytes from adult *Shoc1* KI homozygous (*Shoc1*^KI/KI^) mice and littermate controls (*Shoc1*^KI/+^) co-stained with SYCP3 (green) and RPA2 (red). Scale bars, 10 μm.

**(F)** Quantification of RPA2 foci per cell at indicated meiotic stages using two-tailed Student’s t-test; **** *P* < 0.0001; ns, not significant; error bars, mean ± SEM; n, the total number of nuclei analyzed.

**(G)** Representative images of spread spermatocytes from adult *Shoc1* KI homozygous (*Shoc1*^KI/KI^) mice and littermate controls (*Shoc1*^KI/+^) co-stained with SYCP3 (green) and SPATA22 (red). Scale bars, 10 μm.

**(H)** Quantification of SPATA22 foci per cell at indicated meiotic stages using two-tailed Student’s t-test; **** *P* < 0.0001; ns, not significant; error bars, mean ± SEM; n, the total number of nuclei analyzed. E-Zyg, early zygotene; L-Zyg, late zygotene; Pach, pachytene.

**Figure S8. Isolation of pachytene spermatocytes by modified STA-PUT.**

(**A)** Schematic workflow of the STA-PUT density gradient protocol for isolating pachytene spermatocytes from adult *Shoc1* KI homozygous (*Shoc1*^KI/KI^) mice.

**(B)** Representative images of enriched cell fractions co-stained with SYCP3 (red) and γH2AX. Scale bars, 10 μm.

(**C)** Quantification of germ cell type distributions across fractions from adult *Shoc1*^KI/KI^ male mice; mean ± SEM.

**Figure S9. *Shoc1* KI induced excessive accumulation of DNA damage response (DDR) factors and establishment of MSUC on autosomes.**

**(A)** Representative images of spread spermatocytes from adult *Shoc1* KI homozygous (*Shoc1*^KI/KI^) mice and littermate controls (*Shoc1*^KI/+^) co-stained with SYCP3 (green) and POL II (red). Scale bars, 10 μm.

**(B)** Quantification of POL II foci per cell at indicated meiotic stages using two-tailed Student’s t-test; **** *P* < 0.0001; ns, not significant; error bars, mean ± SEM; n, the total number of nuclei analyzed.

**(C)** Representative images of spread spermatocytes from adult *Shoc1* KI homozygous (*Shoc1*^KI/KI^) mice and littermate controls (*Shoc1*^KI/+^) co-stained with SYCP3 (green) and MDC1 (red). Scale bars, 10 μm.

**(D)** Quantification of MDC1 foci per cell at indicated meiotic stages using two-tailed Student’s t-test; **** *P* < 0.0001; ns, not significant; error bars, mean ± SEM; n, the total number of nuclei analyzed.
