## Supplementary tables for "The XPF-like domain in SHOC1 required for homologous recombination and safeguarding autosome from meiotic silencing of unsynapsed chromatin"

**Supplementary table S1. Information of the primers.**

| **Information** | **Name** | **Sequence (5'-3')** |
| --- | --- | --- |
| Primers for human  gDNA amplification  and Sanger sequencing (5’-3’) | M1: c.A1769G-F | AGGCAGGAGAATAGCTTGAATC |
|  | M1: c.A1769G-R | GGCGAACAGCATCACTTACTA |
|  | M2: c.416_419del-F | CTGCACCCGGCCAATAA |
|  | M2: c.416_419del-R | ATACAGCCAAGGCTGAGAAC |
|  | M3: c.C1582T-F | GACCAAGAACCAGTAAACAGAATAA |
|  | M3: c.C1582T-R | GAGACAGGACTGACAGCTAAA |
|  | M4: c.231_232del-F | CTTCTCTGTGTCTGTTGGAATTTG |
|  | M4: c.231_232del-R | CTTAGAGTTATGCAGGTGGATCTT |
|  | M5: c.1194delA-F | CCAGTGTGGTTCTGTTTATTGTG |
|  | M5: c.1194delA-R | ATGGGACCTAGAGAAATCCAAAG |
|  | M6: c.1347delT-F | CCAGTGTGGTTCTGTTTATTGTG |
|  | M6: c.1347delT-R | AGTCTTTGAGGTGCAAGTCTTAT |
|  | M7: c.G2738-1A-F | ACATTTCAGGTAGAGAATGACCC |
|  | M7: c.G2738-1A-R | ACATTTCAGGTAGAGAATGACCC |
| Primers for genotyping  of *Shoc1* KI mice (5’-3’) | F | GTTATCAGTGGTCACTCCATCCC |
|  | R | AGATTCACTGAGTATCTTCTGGGCT |
| sgRNA for constructing *Shoc1* KI mice (5’-3’) |  | GAGACAGTATGCTTGGCACTGGG |
| Donor oligo for constructing *Shoc1* KI mice (5’-3’) |  | GACACAGCACACTACCCTTTTCACACACACTGATTTCAGTGCTGTTATTTTTCAGACACCCGGTGCCAAGCATACTGTCTCCTAGAAGCCACAGCGAATCCTATCTTAAAGGAGCTTGTTTGC |
| Primers for minigene  of gDNA flanking the  *SHOC1* M7 variant (5’-3’) | F | ATGGATGTTTTCCCAGTCACAGGTATTTATTCTTTTCTTATTGATCTGTAA |
|  | R | GGATATCAGCTGGATGGCAATACTCATTCTCAATATTATCAATGACTCAA |
| Primers for constructing  plasmids for Co-IP (5’-3’) | pCMV-SHOC1-FLAG-F | CTTGCGGCCGGCCGCGAATTCATGACAGATACCTCAGTCTTGGACC |
|  | pCMV-SHOC1-FLAG-R | CCTCTAGAGTCGACTGGTACCTCAAAAAAACCTCAGCCGAGTC |
|  | pcDNA-TEX11-MYC-F | GATTACGCCAAGCTTGGTACCATGGACAATGATGATTTTTTTTCCATGGACT |
|  | pcDNA-TEX11-MYC-R | TGCTGGATATCTGCAGAATTCCTAATCTGACTTGCTCCAGTAGCCA |
|  | pcDNA-C1orf146-MYC-F | GAGGATCTGAAGCTTGGTACCATGGCTGAAAGTGGAAAAGAAAA |
|  | pcDNA-C1orf146-MYC-R | TGCTGGATATCTGCAGAATTCCTAATTTGGGTTAACTGAATCACTATTCA |
|  | pcDNA-M1AP-HA-F | CTTGGTACCGAGCTCGGATCCATGCATCCTGGGCGAACTACT |
|  | pcDNA-M1AP-HA-R | TGCTGGATATCTGCAGAATTCTTAGGGCCTTGAGGGATCCT |
|  | pcDNA-REDIC1-HA-F | CTTGGTACCGAGCTCGGATCCATGAATTGGGTCGGGGGG |
|  | pcDNA-REDIC1-HA-R | TGCTGGATATCTGCAGAATTCTTAGTTGTTCTTATGGATTTCCTGCC |
|  | pCMV-mSHOC1-FLAG-F | CTTGCGGCCGGCCGCGAATTCGAATTCATGGCATTGAACGGA |
|  | pCMV-mSHOC1-FLAG-R | CAGGGATGCCACCCGGGATCCTCTAGATAAAAACTTCAGCCGGG |
|  | pcDNA-mTEX11-HA-F | CTTGGTACCGAGCTCGGATCCATGGACCGCATTACTGACTTTTACT |
|  | pcDNA-mTEX11-HA-R | TGCTGGATATCTGCAGAATTCTTACAGATGGTTTTGAGCTGCCA |
|  | pcDNA-mSPO16-HA-F | CTTGGTACCGAGCTCGGATCCATGGATGAGCGTAGAGGAAAAGA |
|  | pcDNA-mSPO16-HA -R | TGCTGGATATCTGCAGAATTCTCACTCTCCTGAGTCTGCACTAACTG |
|  | pcDNA-mM1AP-HA-F | CAGGGATGCCACCCGGGATCCATGAACCGAAGGAAAACTACTAGTAGAG |
|  | pcDNA-mM1AP-HA-R | CTTGCGGCCGGCCGCGAATTCTTAGGTGTGAGAAGGACGCTCC |
|  | pcDNA-mREDIC1-HA-F | CTTGGTACCGAGCTCGGATCCATGAACTGGGTCGGGGGC |
|  | pcDNA-mREDIC1-HA-R | TGCTGGATATCTGCAGAATTCTTAGAGTGAGTTACTTGTAGGTGTTTCCT |
|  | pCMV-SHOC1-XPF-FLAG-F | CTTGCGGCCGGCCGCGAATTCAAGTGATGCTACAAAAATGCTTTAACA |
|  | pCMV-SHOC1-XPF-FLAG-R | CAGGGATGCCACCCGGGATCCGGAACAGATAGTGTCATTGAAATTCAA |
|  | pCMV-SHOC1-XPF-Mut-FLAG-F | CAGATAGCCGGTGCCAAGCATGTGCCAAGCATTTTGCCTCC |
|  | pCMV-SHOC1-XPF-Mut-FLAG-R | TTGGCACCGGCTATCTGACGCGGCTATCTGACGCTTGAATTTCA |
|  | pCMV-mSHOC1-XPF-FLAG-F | CTTGCGGCCGGCCGCGAATTCAAGTGATGCTGCAAAAATGCTTC |
|  | pCMV-mSHOC1-XPF-FLAG-R | CAGGGATGCCACCCGGGATCCCCAGAGGAGAGATGTGAAGAAAGG |
|  | pCMV-mSHOC1-XPF-Mut-FLAG-F | GTGCCAAGCATACTGTCTCCGTGCCAAGCATACTGTCTCCTAGA |
|  | pCMV-mSHOC1-XPF-Mut-FLAG-R | AGACAGTATGCTTGGCACCGGGTGTCTGAAGCGGGGATCTGGGTGTCTGAAGCGGGGA |
|  | pCMV-SHOC1-FLAG-Venus-N-F | GTGAACCGTCAGAATTAACCATGGACTACAAAGACCATGACGG |
|  | pCMV-SHOC1-FLAG-Venus-N-R | CTCACCCCCCCGGACCCCCCAAAAAACCTCAGCCGAGTCTGC |
|  | pcDNA-TEX11-MYC-Venus-C-F | TAGTCATCGCTATTACCATGGTGATGCGGTTTTGGCAGTAC |
|  | pcDNA-TEX11-MYC-Venus-C-R | TTGTCCCCCCCGGACCCCCCATCTGACTTGCTCCAGTAGCCATG |
|  | pcDNA-TEX11-FLAG-Venus-N-F | AGGATGACGATGACAAGCTTATGGACAATGATGATTTTTTTTCCA |
|  | pcDNA-TEX11-FLAG-Venus-N-R | CTCACCCCCCCGGACCCCCCCTAATCTGACTTGCTCCAGTAGCCA |
|  | pcDNA-C1orf146-MYC-Venus-C-F | TAGTCATCGCTATTACCATGGTGATGCGGTTTTGGCAGTAC |
|  | pcDNA-C1orf146-MYC-Venus-C-R | TTGTCCCCCCCGGACCCCCCATTTGGGTTAACTGAATCACTATTCAG |

**Supplementary table S2. Information of the antibodies.**

| **Name of antibodies** | **Company** | **Catalog**  **number** | **Host** | **Dilution** |
| --- | --- | --- | --- | --- |
| SYCP3 | R&D | AF3750 | Goat | IF (1:25) for human |
| SYCP3 | Abcam | ab97672 | Mouse | IF (1:200) for mice |
| SYCP3 | Abcam | ab15093 | Rabbit | IF (1:200) for mice |
| SYCP1 | Gift from Liu Lab | / | Guinea pig | IF (1:200) |
| γH2AX | Millipore | 2668445 | Mouse | IF (1:500) |
| γH2AX | CST | 9718 | Rabbit | IF (1:500) |
| DMC1 | Santa Cruz | sc-373862 | Mouse | IF (1:200) for human |
| DMC1 | Customed | / | Rabbit | IF (1:100) for mice |
| TRA98 | Abcam | ab82527 | Rat | IF (1:200) |
| pHH3 | CST | 9701 | Rabbit | IF (1:200) |
| c-PARP | CST | 9548 | Mouse | IF (1:200) |
| MLH1 | BD | 51-1327GR | Mouse | IF (1:50) |
| H1T | Proteintech | 18188-1-AP | Rabbit | IF (1:100) |
| HORMAD1 | Proteintech | 13917-1-AP | Rabbit | IF (1:200) |
| SPATA22 | Proteintech | 16989-1-AP | Rabbit | IF (1:100) |
| RPA2 | CST | 2208 | Rat | IF (1:100) |
| RAD51 | Customed | / | Rabbit | IF (1:100) |
| TEX11 | Gift from Yu Lab | / | Goat | IF (1:50) |
| MSH4 | Abcam | Ab58666 | Rabbit | IF (1:50) |
| REDIC1 | Gift from Shi Lab | / | Rabbit | IF (1:100) |
| M1AP | Gift from Shi Lab | / | Rabbit | IF (1:100) |
| HEI10 | Gift from Liu Lab | / | Rabbit | IF (1:100) |
| POL II | Abcam | Ab5095 | Rabbit | IF (1:100) |
| MDC1 | Proteintech | 24721-1-AP | Rabbit | IF (1:200) |
| FLAG-tag | Sigma | F1804 | Mouse | WB (1:5000) |
| HA-tag | CST | C29F4 | Rabbit | WB (1:2000) |
| MYC-tag | CST | 2278 | Rabbit | WB (1:2000) |
| β-ACTIN | Proteintech | 60008 | Mouse | WB (1:5000) |
| Alexa Fluor® 488 Donkey anti-Mouse IgG (H+L) | Invitrogen | A-21202 | Donkey | IF (1:400) |
| Alexa Fluor® 594 Donkey anti-Mouse IgG (H+L) | Invitrogen | A-21203 | Donkey | IF (1: 400) |
| Alexa Fluor® 488 Donkey anti-Rabbit IgG (H+L) | Invitrogen | A-21206 | Donkey | IF (1: 400) |
| Alexa Fluor® 594 Donkey anti-Rabbit IgG (H+L) | Invitrogen | A-21207 | Donkey | IF (1: 400) |
| Alexa Fluor® 488 Donkey anti-Rat IgG (H+L) | Invitrogen | A-21208 | Donkey | IF (1: 400) |
| Alexa Fluor® 488 Donkey anti-Goat IgG (H+L) | Invitrogen | A-11055 | Donkey | IF (1: 400) |

**Supplementary table S3. Clinical and semen characteristics in Chinese men with bi-allelic *SHOC1* variants**

|  | **Subject** | | | | |  |
| --- | --- | --- | --- | --- | --- | --- |
|  | P1 | P2 | P3 | P4 | P5 | Reference |
| Characteristics | | | | | |  |
| Age (years) | 29 | 30 | 25 | 34 | 33 | / |
| Testicular Volume (Left) (ml) | 10 | 15 | 10 | 10 | 15 | 12~15 |
| Testicular Volume (Right) (ml) | 10 | 20 | 10 | 10 | 20 | 12~15 |
| FSH (IU/L) | 12.44 | 3.9 | 4.83 | 3.32 | 5.2 | 1.27~19.26 |
| LH (IU/L) | 5.76 | 5.33 | 4.47 | 3.04 | 4.7 | 1.24~8.62 |
| T (μg/L) | 13.67 | 8.5 | 27.24 | 11.53 | 3.96 | 1.75~7.81 |
| Karyotype | 46,XY | 46,XY | 46,XY | 46,XY | 46,XY | 46,XY |
| Y Chromosome Microdeletions | N | N | N | N | N | N |
| Semen parameters | | | | |  |  |
| Semen volume (ml) |  |  |  |  |  | ≥1.5 |
| Sperm concentration (10^6^/ml) | 0 | 0 | 0 | 0 | 0 | ≥15 |
| PR (%) | 0 | 0 | 0 | 0 | 0 | ≥32 |
| NP (%) | 0 | 0 | 0 | 0 | 0 | / |
| IM (%) | 0 | 0 | 0 | 0 | 0 | / |
| Centrifuged spermatozoa number (/ejaculate) | 0 | 0 | 0 | 0 | 0 | / |

**Abbreviations:** FSH, follicle-stimulating hormone; LH, luteinizing hormone; T, testosterone; PR, progressive; NP, non-progressive; IM, immobility; N, normal phenotype.
